# Actin networks modulate heterogenous NF-κB dynamics in response to TNFα

**DOI:** 10.1101/2022.01.19.476961

**Authors:** Francesca Butera, Julia E. Sero, Lucas G. Dent, Chris Bakal

**Affiliations:** Chester Beatty Laboratories, Division of Cancer Biology, Institute of Cancer Research, 237 Fulham Road, London, SW3 6JB, UK; Biology and Biochemistry Department, Bath University, Claverton Down, BA2 7AY, UK

## Abstract

The canonical NF-κB transcription factor RELA is a master regulator of immune and stress responses and is upregulated in PDAC tumours. Here, we characterised previously unknown endogenous RELA-GFP dynamics in PDAC cell lines by live single cell imaging, which revealed rapid, sustained and non-oscillatory nuclear RELA following TNFα stimulation. Using Bayesian analysis of single cell datasets with variation in nuclear RELA, we computationally predicted that RELA heterogeneity in PDAC cell lines is dependent on F-actin dynamics. By RNA-seq, we identified the actin regulators NUAK2 and ARHGAP31 as transcriptionally regulated by RELA. In turn, *NUAK2* or *ARHGAP31* siRNA depletion downregulates TNFα-stimulated RELA nuclear localisation in PDAC cells, establishing a novel negative feedback loop regulating RELA activation by TNFα. We identify an additional actin-independent feedback loop involving RELB, which suppresses TNFα-mediated RELA nuclear localisation following RELA mediated upregulation of RELB. Taken together, we provide computational and experimental support for interdependence between the F-actin network and RELA translocation dynamics in PDAC.

## Introduction

The NF-κB transcription factor RELA is an essential mediator of the inflammatory and immune responses in all mammals (Hayden et al., 2006) and is central to the canonical NF-κB signalling pathway (Ghosh et al., 1998). As a transcription factor, RELA activation is controlled in large part through its localisation. Inactive RELA is sequestered in the cytoplasm by IκB proteins and IκB degradation by upstream cues, such as the potent inflammatory cytokine Tumour Necrosis Factor α (TNFα), enables RELA translocation to the nucleus where RELA regulates gene expression (DiDonato et al., 1997; Zandi et al., 1997).

Live imaging experiments of fluorescently labelled RELA and electrophoretic mobility shift assays have shown that RELA oscillates between the nucleus and cytoplasm in response to TNFα (Hoffmann et al., 2002; Nelson et al., 2004; Sung et al., 2009; Tay et al., 2010; Sero et al., 2015; Zambrano et al., 2016). Oscillations are driven by a negative feedback loop between RELA and particular IκB isoforms, since the genes encoding IκB proteins are RELA transcriptional targets (Brown et al., 1993; Scott et al., 1993; Sun et al., 1993; Hoffmann et al., 2002). The pattern of RELA translocation has been shown to dictate the specificity and timing of RELA target gene expression, including the genes encoding IκBα, IκBε and the chemokine RANTES (Ashall et al., 2009; Zambrano et al., 2016; Lane et al., 2017). However, most studies characterising RELA translocation dynamics following stimulation use hyperphysiological TNFα doses (e.g. 10 ng/ml) and exogenous RELA reporters.

Previously, we showed that cell shape is a regulator of RELA dynamics in breast cancer cell lines and that breast cancer cells with mesenchymal cell shape (protrusive with low cell-cell contacts) have higher RELA nuclear translocation (Sero et al., 2015). We also identified that RELA activity, coupled to cell shape, is predictive of breast cancer progression (Sailem and Bakal, 2017). Although the mechanistic basis for how cell shape regulates RELA remains poorly understood, studies have shown that chemically inhibiting actin or tubulin dynamics can increase RELA binding to DNA and RELA-dependent gene expression (Rosette and Karin, 1995; Bourgarel-Rey et al., 2001; Németh et al., 2004).

Despite frequent upregulation of both TNFα and RELA in PDAC tumours (Weichert et al., 2007; Zhao et al., 2016), the dynamics and regulation of single cell RELA translocation, as well as RELA transcriptional output, are poorly understood for PDAC cells. Here, we used CRISPR-CAS9 to tag RELA with GFP in the human cell lines MIA PaCa2 and PANC1 and identified rapid, non-oscillatory and heterogenous RELA dynamics by live imaging. To explore potential cytoskeletal or cell shape regulation of RELA in PDAC, we constructed Bayesian models using single cell datasets from TNFα-stimulated PDAC cells with variation in RELA, actin and tubulin measurements for hypothesis generation: one dataset with five PDAC cell lines and another with diverse small-molecules targeting cytoskeletal components. Notably, nuclear RELA was statistically predicted to be dependent on actin features across Bayesian models. Finally, we used RNA-seq to identify genes with distinct patterns of expression based on dependence on TNFα dose and RELA activation. Using live imaging with knockdown of selected targets, our results uncover novel mechanisms regulating inflammation-associated RELA dynamics in PDAC, including negative feedback loops between RELA and the actin modulators NUAK2 and ARHGAP31, and between RELA and the non-canonical NF-κB factor RELB.

## Results

### Single cell endogenous RELA responses to TNFα are non-oscillatory and sustained in PDAC cells

To study PDAC biology, we used the frequently studied human PDAC cell lines MIA PaCa2 and PANC1, which harbour key genomic alterations in PDAC, including KRAS and p53 mutations and homozygous deletions in CDKN2A/p16 (Deer et al., 2010). In addition, MIA PaCa2 and PANC1 cells are epithelial in origin but have distinct cell morphologies and are therefore useful for analysis of cell shape interaction with RELA. To study dynamic RELA localisation changes, we used CRISPR-CAS9 gene editing to fluorescently tag endogenous RELA at the C-terminus with eGFP (abbreviated as RELA-GFP) (Figure 1A and 1B). The C-terminus was selected to avoid interference with the N-terminal Rel homology domain, which contains the transactivation, nuclear import and DNA-binding domains (Kieran et al., 1990; Nolan et al., 1991). GFP+ clones were selected using FACS, single cell sorted and expanded into monoclonal cell lines. We also introduced mScarlet-I to the C-terminus of PCNA (Proliferating Cell Nuclear Antigen; abbreviated as PCNA-Scarlet) – a processivity factor for DNA polymerase δ that functions during replication – which served as a nuclear marker for segmentation and as a cell cycle marker (Kurki et al., 1986; Barr et al., 2017) (Figure 1A and 1B).

**Figure 1:**
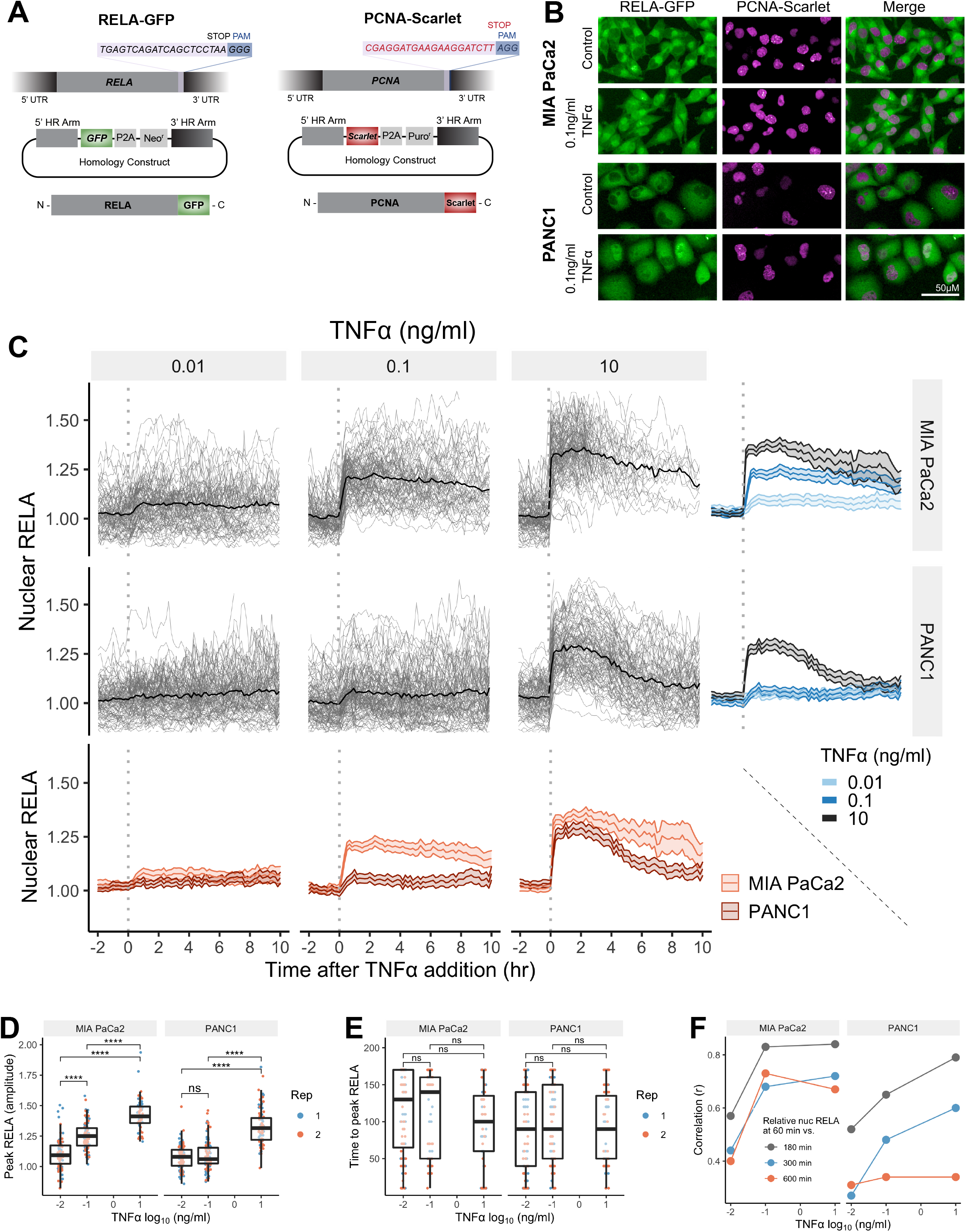
Single cell analysis of live endogenous RELA-GFP dynamics with TNFα in human pancreatic cell lines. (A) Schematic of CRISPR gRNA and homology construct with neomycin (neo) resistance for tagging of the RELA C-terminus with GFP. (B) Confocal microscopy images of MIA PaCa2 and PANC1 monoclonal cell lines expressing endogenously tagged RELA-GFP and PCNA-Scarlet and treated with 1 hr solvent control or 0.1 ng/ml TNFα. (C) Tracks of single cell nuclear RELA-GFP intensity measurements in MIA PaCa2 and PANC1 cells from −120 min to +600 min relative to TNFα addition (0.01 ng/ml, 0.1 ng/ml and 10 ng/ml). n = 50-60 tracked cells per TNFα dose for each of two experimental repeats. (D) Amplitude and (E) time of first nuclear RELA-GFP intensity peak in MIA PaCa2 cand PANC1 single cells. Boxplots show median and interquartile range. M = median per cluster. σ = standard deviation.(F) Pearson’s correlation coefficient (r) between nuclear RELA-GFP intensity at 60 min versus 180, 300 or 600 min within single cell RELA-GFP tracks. Data are normalised within each track to nuclear RELA-GFP intensity at 0 min.

To observe live RELA translocation dynamics in response to inflammatory stimuli, we used timelapse confocal microscopy with automated image analysis to track changes in RELA-GFP localisation on a single cell level in response to TNFα (0.01, 0.1 and 10 ng/ml), from −120 min to +600 min relative to TNFα addition. 0.01 ng/ml TNFα is a physiological dose relevant to healthy and malignant tissue, while 0.1 ng/ml TNFα is detected in highly inflammatory PDAC microenvironments (Zhao et al., 2016). 10 ng/ml TNFα was used in several studies assaying RELA translocation (Hoffmann et al., 2002; Tay et al., 2010; Sero et al., 2015) and is included for comparison, but is substantially above physiological levels (Zhao et al., 2016).

Broadly, we observed a lack of oscillatory nuclear RELA dynamics in PDAC cells, in addition to general maintenance of nuclear RELA across the 10 hr imaging period. MIA PaCa2 cells responded with higher nuclear RELA compared to PANC1 cells at each TNFα dose (Figure 1C). The highest nuclear RELA measurement attained on the single cell level, referred to as ‘peak RELA’, was statistically significantly different between 0.01 and 0.1 ng/ml for MIA PaCa2 but not for PANC1 cells, suggesting that MIA PaCa2 cells are more sensitive to TNFα in terms of RELA activation (Figure 1D). 10 ng/ml TNFα elicited a significantly higher nuclear RELA localisation amplitude in both cell lines compared to lower TNFα doses (0.01 and 0.1 ng/ml). In contrast, TNFα dose did not affect the time to peak RELA in either cell line (Figure 1E), although the time to peak RELA was highly heterogenous on the single cell level within each TNFα dose, suggesting that the time to peak RELA is controlled by multiple cell intrinsic factors.

We also compared RELA measurements within each single cell track to assess the stability of nuclear RELA levels (Figure 1F). In MIA PaCa2 cells, there is a high correlation between nuclear RELA at latter timepoints (180, 300 or 600 min) compared to an early timepoint (60 min) following TNFα addition. PANC1 cells have high correlation between nuclear RELA measurements at 60 min RELA and 180 min, while there is weak correlation between RELA levels at 60 min and 600 min. Thus, in MIA PaCa2, but not PANC1, RELA translocation at 60 min is largely predictive of long-term response to TNFα.

Overall, our data show that RELA nuclear translocation occurs rapidly, and in a non-oscillatory fashion in both MIA PaCa2 and PANC1 cells in response to TNFα. In MIA PaCa2 cells, RELA translocation is largely sustained for hours, while damped in PANC1 cells.

### RELA translocation responses to TNFα are cell cycle independent

To identify whether cell cycle progression contributes to heterogeneity in RELA responses to TNFα in PDAC cells, we categorised each tracked cell by cell cycle stage at the time of TNFα addition, using changes in the appearance and intensity of endogenous PCNA-Scarlet to mark cell cycle transitions (Figure 1 - Supplement 1A). For each cell, we calculated the amplitude and timing of peak nuclear RELA, in addition to the mean nuclear RELA intensity across all timepoints. Largely, there were no differences between cells by cell cycle stage in terms of peak RELA measurements (amplitude or timing) in MIA PaCa2 and PANC1 cells (Figure 1 - Supplement 1C, D). However, we identified statistically significant higher (whole track) mean nuclear RELA in G2 cells in the PANC1 cell line alone (Figure 1 - Supplement 1E).

### TNFα-mediated RELA heterogeneity in PDAC cells is predicted to be dependent on actin dynamics

Because the same TNFα concentration can lead to variable responses, we proposed that there are cell-intrinsic mechanisms that dictate the extent of RELA translocation in PDAC cells. Having previously identified relationships between cell shape and RELA localisation in breast cells (Sero et al., 2015), we hypothesised that differences in actin and tubulin organisation, which regulate cell shape (Machesky and Hall, 1997, Desai and Mitchison, 1997), may explain differences in RELA dynamics. To test this, we expanded our dataset to include immunofluorescence images of the human immortalised PDAC cell lines MIA PaCa2, PANC1, Capan1, SW1990 and PANC05.04. We treated cells with TNFα (1 hr), or with solvent control, and stained for DNA, RELA, F-actin and α-tubulin (Figure 2 - Supplement 1A). We used automated image analysis to segment cell regions and measured 35 geometric, cytoskeletal and Hoechst features, as well as nuclear RELA, in approximately 130,000 cells. The 35 cell features were then reduced by hierarchical clustering to a subset of ten features (Figure 2 - Supplement 1B). Selected features include classical measurements of cell shape (‘Cell Area’, ‘Cell Roundness’ and ‘Nucleus Roundness’), mean measurements and texture analysis of actin and tubulin intensity in the cytoplasm, and the ratio of ‘Actin filament area’ to ‘Cell area’ which assays actin stress fibre abundance (Figure 2 - Supplement 1C).

We used principal component analysis (PCA) to assess the morphological diversity of the five PDAC cell lines using normalised data for cells under control conditions (Figure 2 - Supplement 2). PCA largely clustered data by cell line, indicating distinct cell morphology and cytoskeletal organisation between PDAC cell lines. PCA also validated that the reduced cell feature set is sufficient to capture the morphological heterogeneity in the PDAC lines. Notably, MIA PaCa2 cells scored low on several cell features due to these cells having particularly small cell area, distorted nuclei (low nucleus roundness) and low actin abundance. Moreover, in MIA Paca2 cells, actin is localised in areas of the cytoplasm distal to the membrane. In contrast, PANC1 cells have a notably high cell and nucleus roundness, as well as high membrane/cyt actin, indicating that actin in PANC1 cells is localised predominantly at the cortex.

Distributions of nuclear RELA in the five PDAC cell lines by immunofluorescence revealed highly heterogeneous RELA responses within and across the PDAC cell lines (Figure 2A). To identify features that predict RELA localisation differences, we collated and incorporated normalised single cell measurements across all PDAC cell lines and TNFα treatments into Bayesian networks, harnessing the observed variation in RELA (Figure 2B). Bayesian network models apply statistical inference to heterogeneous experimental data to predict the conditional dependence of components on each other. Bayesian network models appear as influence diagrams consisting of nodes, each representing a measured feature, and arcs that depict predicted dependencies between the nodes. These dependencies represent linear and non-linear relationships, direct and indirect interactions, and illustrate multiple interacting nodes simultaneously (Sachs et al., 2005). We employed a hybrid class of Bayesian algorithm (‘rsmax2’) that generates models with unidirectional arcs using a combination of constraint-based and score-based approaches (Scutari et al., 2018).

**Figure 2:**
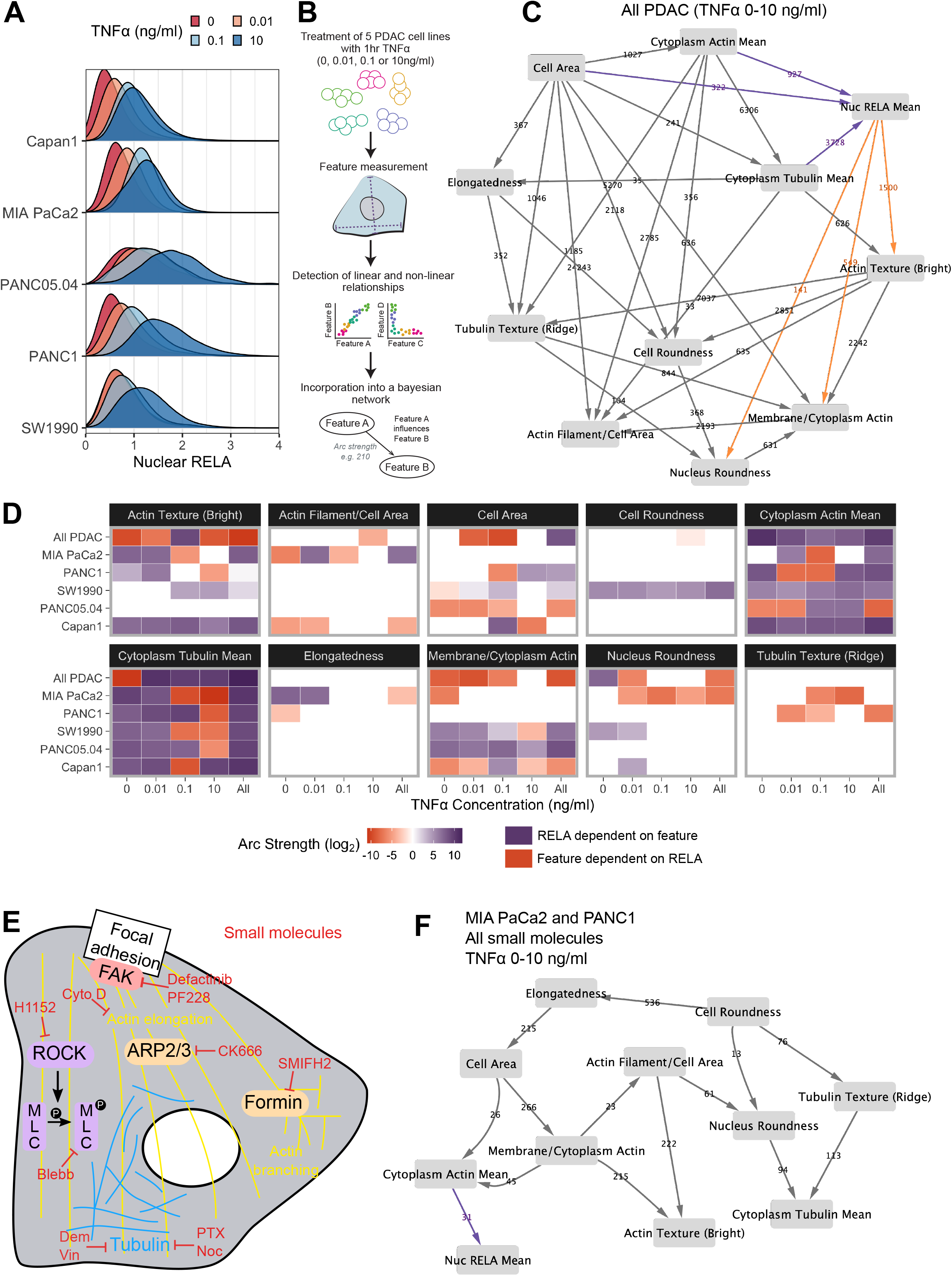
Bayesian network analysis predicts statistical dependence of RELA on actin. (A) Single cell nuclear RELA intensity distributions by immunofluorescence and automated image analysis. (B) Schematic depicting generation of a Bayesian network model for probabilistic relationships between cell features in PDAC cells. PDAC cells were treated with 0, 0.01, 0.1 or 10 ng/ml TNFα for 1 hr. Using automated image analysis of cell markers, features are measured on a single cell basis and Bayesian analysis is used to detect linear and non-linear relationships between the features. These relationships are then incorporated into a Bayesian network model, which is an influence diagram consisting of nodes, each representing a measured feature, and arcs between the nodes that depict predicted dependencies between the nodes. (C) Bayesian network model incorporating data from all PDAC lines treated for 1 hr with 0 (solvent control), 0.01,0.1 or 10 ng/ml TNFα. Values next to arcs represent the strength of the probabilistic relationship expressed by the arc (arc strength). Orange arcs connect features predicted to depend on nuclear RELA mean, and orange arcs connect features predicted to influence nuclear RELA mean. (D) Dependencies involving nuclear RELA mean in Bayesian network models generated with single cell data for individual treatments or cell lines, or for all cell lines collated (top row in each cell feature section), or all treatments collated (rightmost column in each cell feature section). Purple indicates that nuclear RELA mean is predicted to depend on the cell feature in the Bayesian network model. Orange represents that a cell feature is predicted to depend on nuclear RELA intensity. Dependency strengths are calculated as log_2_(|arc strength|), multiplied by −1 for dependencies of cell features on nuclear RELA intensity. (E) Schematic indicating small molecules targeting the cytoskeleton. CK666 inhibits the ARP2/3 complex that mediates actin filament nucleation and branching (Mullins et al., 1998). SMIFH2 inhibits formins (Rizvi et al., 2009), which produce long straight filaments by promoting actin nucleation and filament elongation (Pruyne et al., 2002). Cytochalasin D binds to the growing end of actin filaments and inhibits polymerisation (Schliwa, 1982). H1152 targets Rho-kinase (ROCK), preventing ROCK phosphorylation of myosin light chain that normally promotes actin-binding and contractility, while blebbistatin blocks myosin II ATPase and actin contractility (Sasaki et al., 2002; Kovács et al., 2004). Tubulin-targeting drugs prevent MT assembly (vinblastine and nocodazole), limit MT formation and cause MT depolymerisation (demecolcine), or stabilise MTs and prevent disassembly (paclitaxel) (Spencer and Faulds, 1994; Vasquez et al., 1997; Gigant et al., 2005). Focal adhesion kinase (FAK) regulates turnover of focal adhesions, which are integrin-containing complexes linking intracellular actin to extracellular substrates. (F) Bayesian network model generated by single cell data from MIA PaCa2 and PANC1 cells treated separately with the small molecules in (E) for 3 hr then treated with TNFα (0, 0.01,0.1 or 10 ng/ml) for 1 hr. Numbers indicate arcs strengths. In the presence of small molecule inhibition of the cytoskeleton, nuclear RELA mean is predicted to be dependent on cytoplasm actin mean alone (indicated by the purple arc connecting ‘cytoplasm actin mean’ and ‘nuc RELA mean’).

In order to provide the greatest heterogeneity in nuclear RELA and cytoskeletal/cell shape features for Bayesian model generation, we collated data from all five PDAC lines and TNFα doses (0, 0.01, 0.1 and 10 ng/ml), shown in Figure 2C. This model indicated that nuclear RELA measurements are correlated to and predicted to be dependent on cytoplasmic actin and tubulin intensity, suggesting that cytoskeletal dynamics influence heterogeneity in nuclear RELA translocation with TNFα. Nuclear RELA is also predicted to be dependent on cell area, although with a lower strength of the probabilistic relationship compared to actin/tubulin, while nucleus roundness is predicted to be dependent on nuclear RELA. Interestingly, both actin texture and the ratio of membrane to cytoplasm actin are also predicted to be dependent on RELA, suggesting that RELA and actin dynamics are interdependent. Altogether, our data suggest that the influence of cell shape on RELA translocation we have previously described in breast cancer cells (Sero et al., 2015; Sailem and Bakal, 2017) is likely mediated through cytoskeletal changes.

To understand how inter-line differences in cytoskeletal organisation may influence RELA translocation dynamics, we additionally generated Bayesian network models by subsetting data by cell line and TNFα concentration. We summarised dependencies involving nuclear RELA in Figure 2D. In contrast to our prior findings using Bayesian modelling of RELA and cell shape with breast cancer cells in the absence of cytoskeletal measurements (Sero et al., 2015; Sailem and Bakal, 2017), here we found a general lack of dependence of nuclear RELA on cell shape features, but identified strong and consistent dependencies of nuclear RELA on cytoplasm actin and tubulin, as well as actin texture, in several PDAC cell lines and TNFα doses (Figure 2D). These data computationally predict that actin and tubulin abundance and actin distribution within the cell influence RELA nuclear translocation.

When using datasets of sufficient size, deriving Bayesian models based on largely stochastic and relatively small fluctuations in variables can provide insight into the influence of cytoskeletal components, shape, and RELA on each other (Figure 2C). But molecular interventions provide a means to further drive ordering of connections by identifying regions in state where normally correlated variables become conditionally independent (Pe’er et al., 2001; Sachs et al., 2005). Consequently, we created an additional dataset where cells were imaged following perturbations of tubulin, actin, myosin, or focal adhesion (FA) dynamics (Figure 2E). Such perturbations are intended to alter the value of variables/features (i.e. measures of actin organisation) beyond those observed in normal populations.

For example, CK666 inhibits the ARP2/3 complex – a key mediator of actin filament nucleation and branching (Mullins et al., 1998), while SMIFH2 inhibits formins (Rizvi et al., 2009), which promote actin nucleation and elongation of pre-existing filaments to produce long straight filaments (Pruyne et al., 2002). H1152 targets Rho-kinase (ROCK), preventing ROCK phosphorylation of myosin light chain that normally promotes actin-binding and consequently contractility, while blebbistatin blocks myosin II ATPase and subsequently interferes with actin contractility (Sasaki et al., 2002; Kovács et al., 2004). Tubulin-targeting drugs are commonly used in cancer chemotherapy and either prevent MT assembly (vinblastine and nocodazole), limit MT formation and cause MT depolymerisation (demecolcine), or stabilise MTs and prevent disassembly (paclitaxel) (Spencer and Faulds, 1994; Vasquez et al., 1997; Gigant et al., 2005; Tangutur et al., 2017). We ascertained optimal doses by treating MIA PaCa2 cells with dose ranges for 24 hr, or 3 hr for SMIFH2 (Figure 2 - Supplement 3).

We generated a single cell dataset for use in Bayesian modelling by treating MIA PaCa2 and PANC1 cells with selected drug doses for 2 hr then simultaneously with TNFα (0, 0.01,0.1 and 10 ng/ml) for 1 hr and input these data into the same Bayesian algorithm as above (rsmax2) (Figure 2F). Interestingly, the Bayesian network following perturbations revealed that 5/6 of the variables observed to correlate with RELA (whether they influenced RELA or were influenced by RELA) in untreated cells were independent of RELA in the drug-treated network. Only ‘Cytoplasmic Actin Mean’ remained as a influencing variable following drug-treatment, suggesting that actin abundance is a key regulator of RELA nuclear localisation. While other variables (i.e. Tubulin and cell shape) can indirectly influence RELA, they do so via regulating actin network organisation.

### TNFα dose and duration determine profiles of RELA-dependent gene expression in PDAC cells

As RELA is known to be involved in feedback loops with transcriptional targets, with the best studied example being IκBα, we sought to identify what the transcriptional targets of RELA are in PDAC and which influence RELA in actin dependent and independent ways.

To this end, we carried out RNA-seq analysis of PDAC cells at an early (1 hr) and late (5 hr) timepoint with varying TNFα doses (0.01, 0.1 or 10 ng/ml TNFα) using MIA PaCa2 and PANC1 cells. TNFα treatments were in the presence or absence of dox-induction of IκB super repressor (IκB-SR) to determine which genes require RELA nuclear translocation. 236 genes were differentially expressed in high TNFα (0.1 or 10 ng/ml) versus basal conditions, with the majority of genes (186) upregulated by TNFα (Figure 3A and 3B).

**Figure 3:**
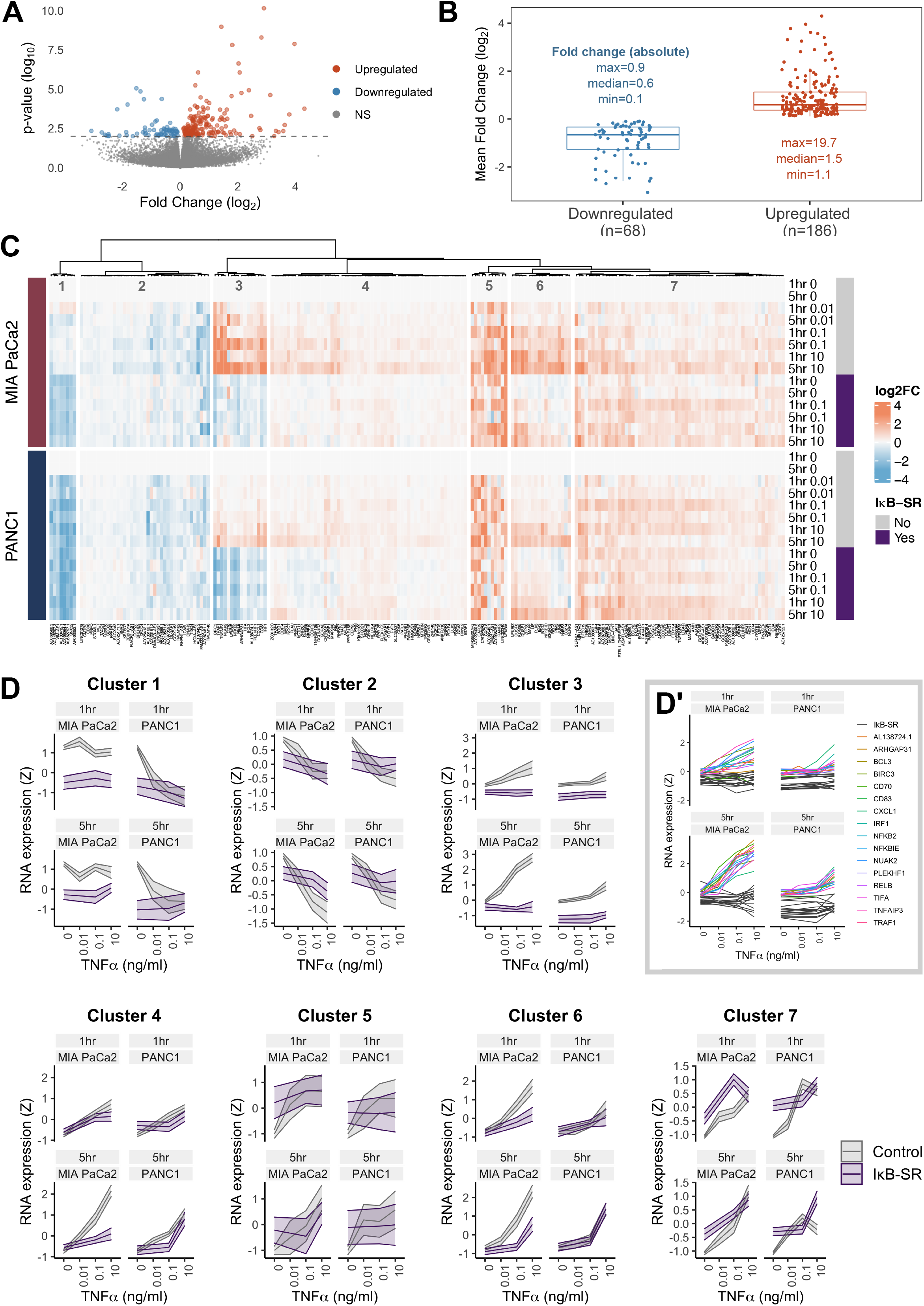
RNA-seq analysis of genes regulated by TNFα and RELA in human PDAC cells. (A) Volcano plot of p-value against mean fold-change (log_2_) per gene comparing RNA expression with high TNFα (0.1 ng/ml or 10 ng/ml) to control conditions (no TNFα) across MIA PaCa2 and PANC1 cells. Counts were normalised and log_2_ fold-changes were calculated using DESeq2. (B) Mean fold-change (log_2_) across in MIA PaCa2 and PANC1 cells for genes significantly downregulated (n = 68) or upregulated (n = 186) by TNFα (p < 0.01) from (A). Also displayed are the maximum, median and minimum absolute fold changes for downregulated and upregulated genes. (C) Clustered heatmap of TNFα regulated genes. Normalised counts from DESeq2 were log_2_ transformed and relative to the respective control (no TNFα or IκB-SR) per timepoint (1 hr or 5 hr) and cell line (MIA PaCa2 or PANC1), then z-scored across all samples independent per gene. Columns are annotated by gene and rows are annotated by cell line, TNFα dose and time, and presence of IκB-SR. (D) Z-scores for all genes per cluster, faceted by cell line and treatment time (1 hr or 5 hr). Ribbons show the 95 % confidence interval and the middle line depicts the mean. Colour corresponds to presence of IκB-SR. (D’) Individual genes within Cluster 3, coloured by gene for control (no IκB-SR) data and grey for all data with IκB-SR (all genes).

To identify patterns of gene expression across TNFα doses and durations, the 236 TNFα-regulated genes were organised by hierarchical clustering into seven clusters, each with distinct expression dynamics or dependence on RELA (Figure 3C and 3D). Cluster 3 shows genes with clear dependence on RELA activation in both MIA PaCa2 and PANC1 cells and has genes significantly upregulated by TNFα in a dose-dependent manner. Moreover, Cluster 3 genes are more highly upregulated by TNFα in MIA PaCa2 compared to PANC1 cells, which is consistent with the RELA dynamics observed by live imaging whereby 0.1 ng/ml is sufficient to increase RELA nuclear localisation in MIA PaCa2 but not PANC1 cells (Figure 1D). Cluster 3 genes also have higher RELA expression at 5 hr TNFα compared to 1 hr TNFα in both cell lines, indicating that they are associated with prolonged RELA activation. As expected from the RELA and TNFα dose dependent properties of this cluster, known RELA targets such as RELB, NFKB2 and BIRC3 are present (Lombardi et al., 1995; Bren et al., 2001; Frasor et al., 2009), and also has the immune ligands CD70 and CD83. Notably, Cluster 3 contains Rho GTPase activating protein 31 (ARHGAP31) and NUAK family kinase 2 (NUAK2), which are involved in the regulation of actin (Tcherkezian et al., 2006; Vallenius et al., 2011).

Two clusters of TNFα-regulated genes in PDAC cells are downregulated by TNFα (Figure 3C and 3D) in a dose-independent manner: Cluster 1 and Cluster 2. Cluster 1 consists of long non-coding RNAs (lncRNAs) and are further downregulated by IκB-SR in the MIA PaCa2 cell line only, suggesting that RELA and TNFα have antagonistic effects on the expression of these lncRNAs. In contrast, Cluster 2 genes are entirely independent of RELA.

The remaining clusters (Figure 3C and 3D) contain genes upregulated by TNFα. Cluster 4 genes are similar to Cluster 3 in that their expression is higher at 5 hr versus 1 hr, but Cluster 4 genes have more moderate upregulation by TNFα. Moreover, Cluster 4 genes appear only to be RELA-dependent with 5 hr TNFα. Clusters 5 and 7 are upregulated by TNFα in a dose independent manner. Finally, genes in Cluster 6 are dependent on TNFα dose but not duration.

Overall, we identified gene expression patterns linked to TNFα dose and duration, as well as variable dependence on RELA, suggesting that TNFα/RELA dynamics determine transcriptional output in PDAC cells.

### RELA modulates the expression of and physically interacts with non-canonical NF-κB and the IκB proteins in PDAC cells

Having observed sustained and non-oscillatory nuclear RELA with TNFα (Figure 1), we decided to consider in more detail how the expression of NF-κB transcription factors and IκB family proteins are affected by TNFα dose and RELA inactivation by IκB-SR in PDAC cells, given that IκB proteins are known regulators of RELA across cell types (Baeuerle and Baltimore, 1988).

RNA-seq in MIA PaCa2 and PANC1 cells revealed that expression of *RELA* itself did not significantly scale with TNFα dose, while *RELB* and the transcriptionally incompetent NF-κB proteins *NFKB1* and *NFKB2* showed increasing expression with increasing TNFα (Left graph in Figure 3 - Supplement 1A). TNFα addition significantly altered the expression of a subset of IκB protein-encoding genes, which display distinct expression dynamics: *NFKBIA* (Figure 3: Cluster 5), *NFKBIB* (Figure 3: Cluster 4), *NFKBIE* (Figure 3: Cluster 3), *NFKBID* (Figure 3: Cluster 6) and *NFKBIZ* (Figure 3: Cluster 6). Of these, *NFKBIA* has the highest absolute RNA expression in MIA PaCa2 cells and *NFKBIB* has the highest abundance in PANC1 cells, while *NFKBID* and *NFKBIZ* have the lowest RNA expression in both cell lines (right graph in Figure 3 - Supplement 1A).

T-test comparison with multiple test correction of RNA expression in TNFα-treated cells revealed that *RELB* and *REL* have significantly reduced expression with IκB-SR presence versus absence in both MIA PaCa2 and PANC1 cells, while *NFKB2* is only affected by IκB-SR induction in PANC1 cells (Figure 3 - Supplement 1A’). Furthermore, expression of the canonical RELA binding partner *NFKB1* is unaffected by RELA-inactivation by IκB-SR in both cell lines. Similarly, expression of the IκB family protein-encoding genes *NFKBIA* and *NFKBIB* were not significantly impacted by RELA inactivation in either cell line, while *NFKBIE* expression was reduced with IκB-SR in PANC1. Therefore, of the core NF-κB signalling components, *RELB*, *REL* and *NFKBIE* expression appear to be dependent on TNFα-stimulated RELA nuclear translocation in PDAC cells.

We also considered how expression of NF-κB and IκB genes are affected by the treatment duration of TNFα (Figure 3 - Supplement 1A’’), combining data from 0.1 ng/ml and 10ng/ml TNFα. In both MIA PaCa2 and PANC1 cells, *RELB* expression was significantly higher with 5 hr compared to 1 hr TNFα, which was also the case for *NFKB2* and *NFKBIE* in the PANC1 cell line alone.

Lastly, we checked which NF-κB/IκB proteins interact with RELA in PDAC cells. GFP-Trap followed by mass spectrometry with MIA PaCa2 cells expressing endogenously tagged RELA-GFP pulled down six NF-κB/IκB proteins (Figure 3 - Supplement 1B). IκBα/NFKBIA, IκBβ/NFKBIB and IκBε/NFKBIE were pulled down at lower abundance with increasing TNFα dose, in line with the well-studied degradation of IκB proteins downstream of TNFα stimulation (Chen et al., 1995). Interestingly, the NF-κB protein REL also had reduced interaction with RELA with TNFα. In contrast, NFKB1 and NFKB2 showed increased interaction with RELA with TNFα. RELA therefore appears to form NF-κB heterodimers with REL, NFKB1 and NFKB2 in PDAC cells.

### The NF-κB signalling components RELB and IκBβ and the actin modulators ARHGAP31 and NUAK2 suppress TNFα-stimulated RELA nuclear localisation in PDAC cells

To test for potential feedback loops regulating RELA in PDAC, we evaluated the effect of siRNA knockdown of TNFα-regulated genes on TNFα-stimulated RELA dynamics. We targeted the genes encoding the known RELA inhibitors IκBα, IκBβ and IκBε, which were pulled down with RELA-GFP by GFP-Trap, and the non-canonical NF-κB transcription factors RELB and NFKB2. In addition, we tested knockdown of the Cluster 3 (from Figure 3) genes *NUAK2* and *ARHGAP31*, which are known actin modulators (Tcherkezian et al., 2006; Vallenius et al., 2011). MIA PaCa2 and PANC1 cells were transfected with siRNAs for 48 hr then live imaged for 2 hr prior to until 12 hr following addition of 10 ng/ml TNFα or solvent control (Figure 4A-D). We used non-targeting (NT) siRNA as a negative control for comparison and verified transfection efficacy using RELA siRNA, which abrogated the nuclear RELA signal (Figure 4A).

**Figure 4:**
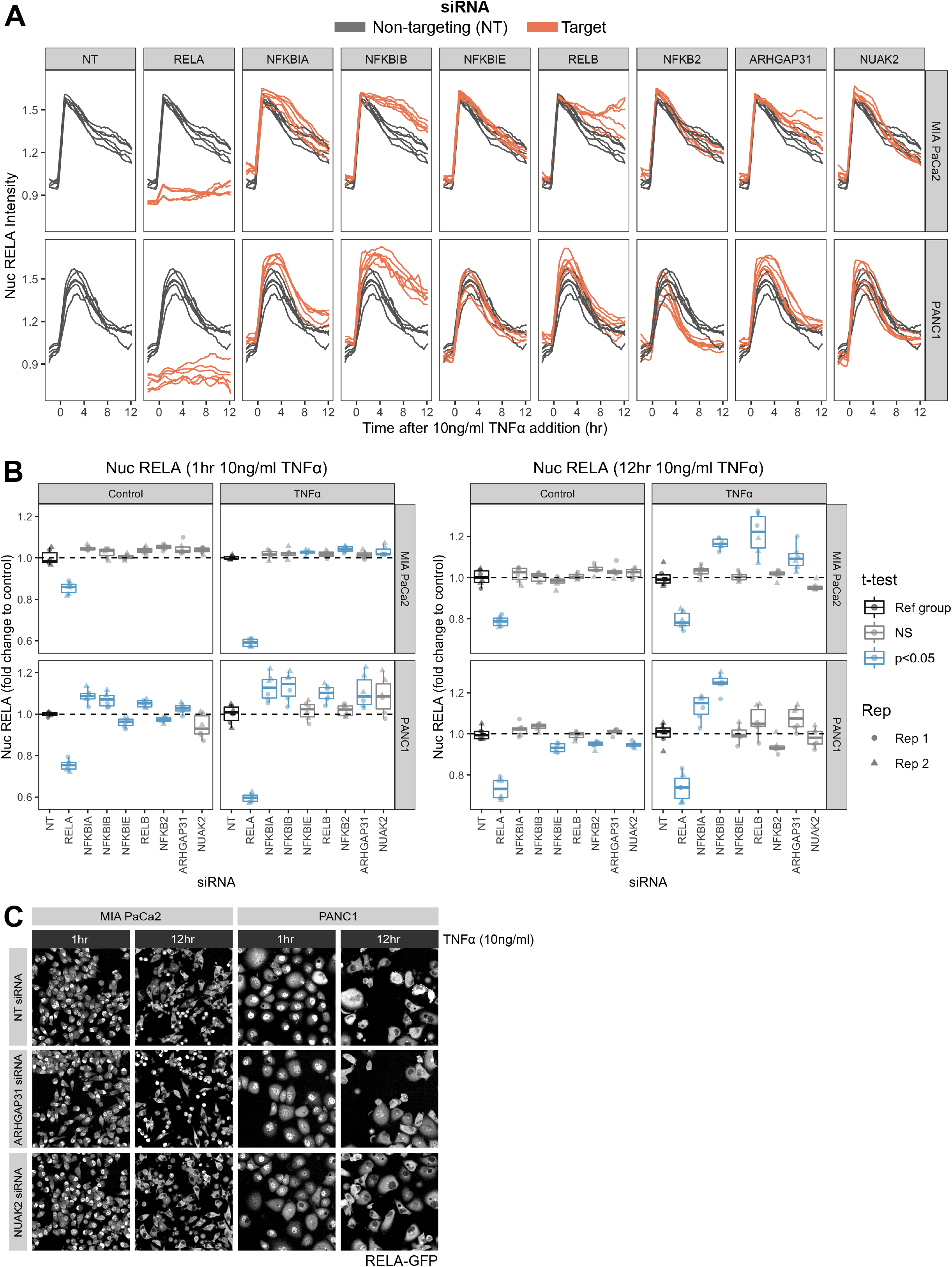
Effect of siRNA knockdown of TNFα and actin signalling on RELA dynamics. (A) Tracks of RELA-GFP intensity measurements relative to 10 ng/ml TNFα addition (−2 hr to +12 hr) in MIA PaCa2 and PANC1 cells following 48 hr siRNA incubation. Data show well means from two experimental repeats. Grey lines represent RELA-GFP measurements with non-targeting (NT) siRNA and are identical in all facets. Orange lines show RELA-GFP tracks from cells with siRNA depletion for the target gene indicated in the column facet. (B) Nuclear RELA-GFP intensity measurements (well means from two experimental repeats) for data in at 1 hr (left graph) or 12 hr (right graph) following 10 ng/ml TNFα addition. Statistical results are shown for multiple t-tests with Benjamini-Hochberg correction comparing measurements from each siRNA to the non-targeting siRNA control, independently by cell line and treatment (0 or 10 ng/ml TNFα). All data are normalised to the mean nuclear RELA intensity of the non-targeting control by experimental repeat (n = 2) and cell line. Blue boxplots represent statistically significant (p < 0.05) results and grey boxplots show statistically non-significant results (p < value). (C) Snapshots from confocal microscopy timelapse imaging of endogenous RELA-GFP and PCNA-Scarlet (nuclear marker) in MIA PaCa2 and PANC1 cells following NT, ARHGAP31 or NUAK2 48 hr siRNA treatment, followed by 1 hr or 12 hr 10 ng/ml TNFα addition.

To identify whether siRNAs affected the early/peak RELA and sustained RELA response to TNFα, we calculated the fold change of mean nuclear RELA with each siRNA to NT siRNA at 1 hr or 12 hr TNFα stimulation, and compared fold changes using t-test with multiple comparison correction (Figure 4B and 4C). Considering TNFα/NF-κB pathway components, we found that several siRNAs enhanced nuclear RELA in PANC1 but not in MIA PaCa2 in basal conditions, including *NFKBIA, NFKBIB, RELB* and *ARHGAP31*, suggesting that RELA activation is actively and constitutively suppressed by several mechanisms in PANC1 cells.

Focusing on the early RELA response (1 hr) to TNFα, we found that *NFKBIE* and *NFKB2* depletion reduced nuclear RELA in PANC1 in control conditions, suggesting that these TNFα/NF-κB signalling components enhance nuclear RELA in basal conditions. Meanwhile, depletion of *NFKBIA, NFKBIE*, and *RELB* significantly increased nuclear RELA in basal and TNFα conditions in PANC1.

In terms of the RELA response to 12 hr post-TNFα, we observed more pronounced effects with TNFα in MIA PaCa2 compared to with 1 hr post-TNFα. In particular, depletion of *RELB* and *NFKBIB*, but not *NFKBIA* or *NFKBIE*, significantly enhanced nuclear RELA. In PANC1 cells, *NFKBIE* and *NUAK2* siRNA depletion reduced nuclear RELA in basal conditions. In contrast, none of these gene depletions affected TNFα-regulated nuclear RELA with PANC1, which was significantly increased by depletion of *NFKBIA* and *NFKBIB*.

Overall, *RELB* and *NFKBIB* suppressed nuclear RELA in multiple contexts and may therefore be key regulators of RELA dynamics with TNFα in PDAC. However, despite implication of IκBβ and IκBε in the suppression of RELA oscillations (Hoffmann et al., 2002), we did not observe RELA oscillations with *NFKBIB* and *NFKBIE* depletion in PDAC cells, suggesting that the lack of RELA oscillations in PDAC cells may be more related to IκBα or via independent positive feedback mechanisms.

We then focused on the effect of siRNA knockdown of the putative actin regulators and TNFα/RELA-dependent genes *ARHGAP31* and *NUAK2* on early (1 hr post-TNFα) and late (12 hr post-TNFα) nuclear RELA in MIA PaCa2 and PANC1 cells, in order to explore the dependence of RELA on actin dynamics computationally predicted by Bayesian modelling (Figure 2). Strikingly, *ARHGAP31* depletion significantly increased the early (1 hr) response of PANC1 cells to TNFα and the late (5 hr) response of MIA PaCa2 to TNFα, while NUAK2 depletion increased nuclear RELA with TNFα in MIA PaCa2 cells (Figure 4B and 4C). Thus, *NUAK2*/*ARHGAP31* expression impacts TNFα-mediated RELA dynamics in PDAC cells.

We additionally checked whether depletion of *ARHGAP31* or *NUAK2* altered actin and cell shape using Phalloidin staining with each siRNA (Figure 4 - Supplement 1). MIA PaCa2 and PANC1 cells in the presence of *ARHGAP31* siRNA showed flatter morphology and reduction of stress fibre abundance, while *NUAK2* siRNA visibly increased actin abundance and the presence of lamellipodia in both cell lines.

Therefore, we identified RELA targets (*ARHGAP31/NUAK2*) involved in actin modulation that additionally downregulate nuclear RELA with TNFα, providing a potential mechanism that may mediate the relationship between actin and RELA.

## Discussion

To characterise previously unknown single cell RELA dynamics in PDAC cells, we used CRISPR-CAS9 genome editing to tag RELA endogenously with GFP in the human PDAC cell lines MIA PaCa2 and PANC1. Using live imaging and automated image analysis, we characterised RELA responses to the cytokine TNFα, which is upregulated with PDAC progression (Zhao et al., 2016). Strikingly, PDAC cells show atypical RELA dynamics compared to reports in cells from other tissues (Ashall et al., 2009; Tay et al., 2010; Sero et al., 2015), as we observed that PDAC cells maintain prolonged nuclear RELA localisation and RELA responses are cell cycle independent. However, we identified that a key difference between MIA PaCa2 and PANC1 cells is that MIA PaCa2 cells respond with higher and sustained RELA nuclear localisation compared to PANC1 cells, while PANC1 cells have a damped RELA response over time. Similar to other cell lines (Ashall et al., 2009; Tay et al., 2010; Sero et al., 2015), we found that RELA nuclear translocation in PDAC cells occurs immediately following TNFα addition and peak nuclear RELA localisation occurs around 1 hr post-TNFα.

The distinct RELA dynamics between PDAC with reports from other tissues may be attributed to the use of knock-in RELA-GFP in this study, since most studies used exogenous RELA fusion constructs (Ashall et al., 2009; Tay et al., 2010; Sero et al., 2015). As RELA upregulates several genes involved in positive and negative feedback (Collart et al., 1990; Libermann and Baltimore, 1990; Brown et al., 1993; Scott et al., 1993; Sun et al., 1993), which we observed in the present study using RNA-seq, RELA overexpression could interfere with its intrinsic dynamics. Nonetheless, other studies that have tagged RELA endogenously did detect oscillations, including MEFs (Sung et al., 2009; Zambrano et al., 2016) and MCF7 breast cancer cells (Stewart-Ornstein and Lahav, 2016).

The high sensitivity of PDAC cells to TNFα and lack of oscillations suggest that PDAC cells may suppress negative feedback imposed by IκBα, as Hoffman et al. (2002) demonstrated that oscillations in NF-κB DNA binding are due to negative feedback with IκBα, which is encoded by a gene (*NFKBIA*) upregulated by NF-κB factors. Hoffman et al. also identified that cells with high IκBβ and IκBε expression, or absence of IκBα, lose oscillations stimulated by TNFα. These findings motivated our inspection into RELA interaction with IκB proteins and upregulation of IκB genes by TNFα in PDAC cells. Using co-immunoprecipitation of RELA with MIA PaCa2 cells, we found that RELA binds to IκBα, IκBβ and IκBε and binding to each is suppressed in high TNFα concentrations. However, contrary to our expectations based on the findings by Hoffmann et al., we did not observe RELA oscillations with *NFKBIB* or *NFKBIE* depletion in MIA PaCa2 or PANC1 cells, suggesting that the lack of RELA oscillations in these cell lines may be due to enhanced positive feedback and/or insufficient *NFKBIA* expression.

We used Bayesian modelling as an unbiased and high dimensional approach to determine whether descriptors of cell shape and the cytoskeleton correlate with the observed heterogeneity in RELA localisation, having previously used Bayesian modelling to show that RELA localisation in breast cells is strongly dependent on neighbour contact, cell area, and protrusiveness in the presence and absence of TNFα (Sero et al., 2015). In the present study, we extended the analysis to include measurements of actin and tubulin organisation. Since Bayesian modelling relies on heterogeneity in measurements to make predictions, we used a dataset with five PDAC cell lines with high intra and inter-line variability in RELA, as well as distinct cell shape and cytoskeleton between the cell lines. We independently ran Bayesian modelling with a dataset of cells with perturbation of the cytoskeleton using small molecules, based on the seminal study by Sachs et al. (2005) that used Bayesian inference to derive cell signalling networks. Consistently among our models, differences in cytoplasmic actin intensity, as well as measures of actin localisation (cortical versus cytoplasmic actin), were predictive of differences in nuclear RELA between single PDAC cells.

To identify potential regulatory loops involving RELA in PDAC, we used RNA-seq analysis with MIA PaCa2 and PANC1 cells treated with varying TNFα doses +/- IκB-SR. We identified 236 genes significantly regulated by TNFα, which may be viewed as candidates for targeting NF-κB signalling or output in PDAC. Identifying novel targets for PDAC is important as the 5-year survival rate of non-resected PDAC patients has remained unchanged from 1975 to 2011 (Bengtsson et al., 2020). Moreover, RELA is hyperactive in 50-70 % of PDAC tumours (Wang et al., 1999; Weichert et al., 2007) and contributes to both cancer progression (Fujioka et al., 2003; Melisi et al., 2009) and resistance to chemotherapy (Bold et al., 2001; Kunnumakkara et al., 2007).

Of note, we present the first evidence that *ARHGAP31* is a transcriptional target of TNFα or RELA. Moreover, ARHGAP31 is not previously associated with pancreatic cancer, but has well studied roles in actin modulation, since ARHGAP31 is the human orthologue for the mouse protein CdGAP (mCdc42 GTPase-activating protein) and ARHGAP31 inactivates the GTPases CDC42 and RAC1 (Lamarche-Vane and Hall, 1998; Tcherkezian et al., 2006). CDC42 is involved in the formation of premigratory filopodia in PDAC cells and promotes invasiveness (Razidlo et al., 2018), while RAC1 expression is required for the development of PDAC tumours, in addition to the formation of the ADM and PanN precursors in KRAS^G12D^ mouse models (Heid et al., 2011), indicating that RAC1 may play a role in actin remodelling during PDAC initiation.

Our results also demonstrate that TNFα upregulates RNA expression of the kinase-encoding gene *NUAK2* (SNARK), while *NUAK2* expression is abrogated in the presence of IκB-SR. Our identification of a feedback loop between RELA nuclear translocation and *NUAK2* expression is analogous to prior findings that NUAK2 increases the activity of the mechanosensitive transcriptional coactivator YAP through stimulation of actin polymerisation and myosin activity (Yuan et al., 2018). A potential source for further study could consider myosin phosphatase target subunit 1 (MYPT1), a substrate for NUAK2 (Yamamoto et al., 2008) that is reported to mediate NUAK2 regulation of actin stress fibres in growing cells (Vallenius et al., 2011). From a therapeutic perspective, NUAK2 may be a promising target to follow-up for potential use in PDAC therapy, as kinases contain an ATP-binding cleft that is a druggable pocket and can contain additional druggable sites distal to the ATP or substrate binding pockets conferring inhibitor specificity (Lamba and Ghosh, 2012). However, NUAK2 is a prognostic marker for PDAC and is associated with a favourable prognosis according to The Human Protein Atlas, which proposes a potential tumour suppressor role for NUAK2 in PDAC, suggesting that enhancing NUAK2 may be a favourable strategy for PDAC. Therefore, our data provide incentive for exploration of a RELA-NUAK2 signalling axis in PDAC progression and response to therapy.

## Author Contributions

F.B., J.E.S., and C.B. conceived the study. F.B. performed the experiments and image analysis. F.B. and C.B. wrote the manuscript with support and discussion from J.E.S. and L.D..

## Acknowledgements

We thank Theodoros I Roumeliotis and Jyoti Choudhary for performing protein mass spectrometry and analysis. We thank Andrea Brundin for assistance with single cell tracking. We gratefully acknowledge funding for this work by Cancer Research UK, awarded to F.B. (S_3567).

## Materials and Methods

Further information and requests for resources should be directed to and will be fulfilled by Chris Bakal.

### Cell Lines and Cell Culture

Cell lines were maintained at 37°C and 5% CO2 in Dulbecco’s Modified Eagle Medium (DMEM; Gibco) supplemented with 10% heat-inactivated Fetal Bovine Serum (Sigma) and 1% Penicillin/Streptomycin (Gibco). MIA PaCa-2, PANC-1, Capan-1, SW-1990, and Panc05.04 were obtained from ATCC.

### Generation of cell lines with fluorescently tagged RELA and PCNA by CRISPR-CAS9

RELA and PCNA were tagged endogenously at each C-terminus using CRISPR-CAS9-mediated gene editing in MIA PaCa2 and PANC1 cells. RELA was tagged with enhanced GFP (Zhang et al., 1996) and PCNA was tagged with mScarlet-I (Bindels et al., 2017), abbreviated here as RELA-GFP and PCNA-Scarlet respectively. RELA-GFP was first introduced into wildtype cell lines then PCNA-mScarlet was added to validated RELA-GFP clones.

Homology constructs were generated by extracting the region around the stop codon of each gene by PCR. The product was used as a template to amplify the left homology arm (LHA) and right homology arm (RHA) by PCR. PCRs were carried out using High-Fidelity Q5 DNA Polymerase (NEB) according to the manufacturer’s protocol. The RHA contains a mutation corresponding to the gRNA protospacer adjacent motif (PAM) to prevent repeat targeting by the Cas9 nuclease. Primers used to amplify the homology arms included overlaps for 1) a DNA cassette encoding a linker protein, the fluorescent protein, and antibiotic resistance (kindly donated by Francis Barr); 2) the pBluescript II SK (-) vector (Agilent) following EcoRV digestion. The final homology construct was generated from the four DNA oligos by Gibson assembly using the NEB Gibson Assembly Master Mix and according to the NEB protocol.

gRNA oligos were designed using CRISPR.mit.edu. Forward and reverse oligos were phosphorylated, annealed and ligated into a BbsI-digested pX330 U6 Chimeric hSpCas9 plasmid, gifted from Feng Zhang (Cong et al., 2013).

Custom oligos were synthesised by Sigma-Aldrich. gRNA oligos were designed using CRISPR.mit.edu.: TGAGTCAGATCAGCTCCTAA (RELA gRNA forward), TTAGGAGCTGATCTGACTCA (RELA gRNA reverse), CGAGGATGAAGAAGGATCTT (PCNA gRNA forward), AAGATCCTTCTTCATCCTCG (PCNA gRNA reverse). The following oligos were used for genomic DNA extraction: TGGGTCAGATGGGGTAAGAG (RELA C-terminus forward), CCAGCTTGGCAACAGATTTA (RELA C-terminus reverse), GCCCTGGAGCCTTGATATTCA (PCNA C-terminus forward), TCTCACTTGTTCCTTGAGCTCA (PCNA C-terminus reverse). The following oligos were used for homology construct generation, using the genomic DNA as a template: ACGGTATCGATAAGCTTGATTGGGTCAGATGGGGTAAGAG (RELA left arm forward), CGCCACCACCGCTCCCACCGGAGCTGATCTGACTCAGCA (RELA left arm reverse), TTCTTGACGAGTTCTTCTGAGGAGGTGACGCCTGCCCTCC (RELA right arm forward), CCGGGCTGCAGGAATTCGATCAAGGAAGTCCCAGACCAAA (RELA right arm reverse), ACGGTATCGATAAGCTTGATGAGTTTGCAGAGCTGAAATTA (PCNA left arm forward), CGCCACCACCGCTCCCACCAGATCCTTCTTCATCCTCGA (PCNA left arm reverse), GTTATGTGTGGGAGGGCTAAGCATTCTTAAAATTCAAGAA (PCNA right arm forward), CCGGGCTGCAGGAATTCGATTCTCACTTGTTCCTTGAGCT (PCNA right arm reverse).

Cells were transfected with homology and gRNA constructs using Lipofectamine 2000 (ThermoFisher) according to the manufacturer’s protocol. Cells were expanded and selected for antibiotic resistance for three weeks. FP-positive cells were selected using FACS and sorted into single cells per well in 96-well plates and clones were expanded and tested for FP presence by amplifying and sequencing the C-terminus of the RELA and PCNA genes from genomic DNA, in order to confirm the presence of the linker and eGFP or mScarlet DNA.

### Cell seeding and treatment for fixed image analysis

Cells were seeded at a density of 1,000 cells per well in 384-well plates unless otherwise specified.

For comparison of cell shape and cytoskeletal features in the five PDAC cell lines, cells were fixed 2 days after seeding, including 1 hr TNFα treatment. The experiment was carried out with three times (biological replicates) in total, each with four technical replicates (wells) per condition.

### TNFα treatment

Cells were treated with human recombinant TNFα diluted in complete medium at a final concentration of 0.1 ng/ml, or 1 ng/ml and 10 ng/ml when specified. TNFα sourced from Sino Biological was diluted in water and used to treat the panel of PDAC lines for Bayesian analysis (Figure 2). Due to lack of availability, TNFα was sourced from R&D Systems and diluted in 0.1% BSA/PBS for all other experiments.

### Immunofluorescence

Cells were fixed with warm formaldehyde (FA) dissolved in PBS at a final concentration of 4% for 15 min at 37°C then washed three times with PBS. Cells were permeabilised in 0.2% TritonX-100 (Sigma Aldrich) dissolved in PBS for 10 min and blocked in 2% BSA/PBS for 1 hr at RT (room temperature). Cells were stained with 10μg/ml Hoechst (Sigma Aldrich) in PBS (1:1,000) for 15 minutes, washed three times and left in PBS/azide before imaging.

Cells were incubated with primary antibodies for 2 hr at RT or overnight at 4 °C, washed three times with PBS, and incubated with secondary antibodies for 90 min at RT.

Primary antibodies used were rabbit anti-p65/RELA NF-κB (Abcam #16502; 1:500), rat anti-α-tubulin (Bio-Rad #MCA78G; 1:1000), and rabbit anti-pFAK Tyr397 (Invitrogen #44-624G; 1:250).

Cells were incubated with secondary antibodies for 90 min at RT. Secondary antibodies used were Alexa 647 goat anti-rat IgG (Invitrogen #A21247), Alexa 488 goat anti-rabbit IgG (Invitrogen #A11034), and Alexa 647 goat anti-rabbit IgG (Invitrogen #A21246).

For F-actin staining, cells were incubated with Alexa-568 phalloidin (lnvitrogen #A12380; 1:1000) for 90 min simultaneously with secondary antibodies.

### Imaging and automated analysis of fixed cells

A minimum of 21 fields of view per well were imaged using the PerkinElmer Opera confocal microscope using a 20x air objective. Image analysis was performed using custom image analysis scripts created and executed on PerkinElmer’s Columbus 2.6.0 software platform. Scripts detected and segmented individual nuclei using Hoechst and the cytoplasm using Tubulin, or RELA when Tubulin is not included in the staining set. Cells touching the image border are filtered out and neighbour contact (% cell border touching another cell) for each remaining cell is calculated. The nuclear region is reduced by 1 px from the nuclear outer border from Hoechst segmentation and the ring region is set as the area 2 px to 6 px outside of the nuclear outer border. Intensities of all stains are calculated in all segmented regions on a single-cell level. A total of 32 geometric, cytoskeletal and Hoechst features were measured in addition to measurements of RELA/RELA-GFP. Texture features were calculated using SER methods with region normalisation. Bright and Spot textures were smoothed to a kernel of 4px to detect large patches (bundles) of actin/tubulin. Ridge texture was non-smoothed to detect sharp ridges (filaments) of actin/tubulin. Elongatedness was calculated as ((2*Cell Length)^2^/Cell Area). Actin Filament Area was measured using Columbus’s ‘Find Spots’ function applied to the actin channel. Neighbour contact was calculated using an inbuilt Columbus algorithm calculating the percentage of a cell’s border in contact with other cell borders. Grouped neighbour contact measurements were generated from non-normalised data rounded to the nearest multiple of ten.

### Live cell imaging and analysis

MIA PaCa2 RELA-GFP PCNA-Scarlet and PANC1 RELA-GFP PCNA-Scarlet cells were seeded (1,000 cells/well) in a 384-well plate one day prior to imaging. 4 fields per well were imaged at 10 min intervals with a 20x air objective and an environmental control chamber set to 80% humidity, 5% CO2 and 37°C.

For live imaging with TNFα only (no siRNA), cells were imaged for 2 hr prior to and 48 hr following TNFα addition using the Opera QEHS imaging system (PerkinElmer). Nuclear RELA and PCNA intensities were measured using Nuclitrack software (Cooper et al., 2017), with 50-60 cells tracked per treatment, cell line and biological replicate (n = 2). Cells were tracked for 10 hr following TNFα treatment while the total 48 hr imaging period was used to ascertain cell fate (division or death). Each biological replicate consisted of eight technical (well) replicates per cell line and treatment. Missing data points in the center of tracks were imputed using the ‘imputeTS’ package in R.

For live imaging with TNFα and siRNA, cells were imaged for 2 hr prior to and 12 hr following TNFα addition using the Opera Phenix Plus High-Content Screening System (PerkinElmer), with two biological replicates (n = 2). Nuclear RELA was measured in all non-border cells using Harmony software (PerkinElmer). In R, data were smoothed with a Savitzky-Golay filter using the ‘Signal’ package (filter order p = 1, filter length n = 7).

In all experiments, intensity measurements are normalised to the mean of the solvent control measurements (TNFα absence) by cell line, to account for photobleaching and laser power changes. Data with siRNAs are normalised to the non-targeting siRNA controls (solvent control).

Nuclear RELA peaks were detected in R as the maximum nuclear RELA intensity by track following Savitzky-Golay filtering using the ‘Signal’ package (filter order p = 3, filter length n = 11).

### GFP-Trap

MIA PaCa2 RELA-GFP cells were cultured in 15 cm dishes to 80 % confluence then treated with TNFα at a final concentration of 0.01 ng/ml, 0.1 ng/ml or 10 ng/ml, or with a solvent control. Co-immunoprecipitation (Co-IP) was carried out using GFP-Trap Agarose beads (Chromotek #gta-10) or with binding control agarose beads (Chromotek #bab-20) using RIPA lysis buffer and according to the manufacturer’s protocol. Samples were analysed using mass spectrometry by the ICR proteomics core (Theodoros I Roumeliotis and Jyoti Choudhary).

### Cytoskeletal Drug Treatments

2,000 MIA PaCa2 RELA-GFP cells/well were seeded in 384-well plates and treated the next day with the following drugs for 24 hr without TNFα at the specified dose ranges: Paclitaxel (6.25-200 nM; Sigma), Vinblastine (3.125-100 nM; Sigma), Nocodazole (12.5-400 nM; Sigma), Demecolcine (6.25-200 nM; Sigma), Cytochalasin D (0.125-4 μM; Sigma), CK666 (12.5-400 μM; Sigma), H1152 (1.25-40 μM; Tocris), Blebbistatin (1.25-40 μM; Sigma), PF573228 (0.625-20 μM; Tocris), and Defactinib (0.625-20 μM; Selleckchem). Cells were treated with SMIFH2 (3.125-100 μM; Abcam) for only 3 hr due to reported cycles of de- and re-polymerisation of 4-8 hr and inefficacy after 16 hr (Isogai et al., 2015). Ranges were selected according to literature and manufacturers’ recommendations. Doses for further analysis were selected based on the observed effect on the cytoskeletal target and cell morphology.

MIA PaCa2 and PANC1 RELA-GFP cells were seeded at 2,000 cells/well in 384-well plates and treated the following day with selected doses of the small-molecule inhibitors for 2 hr plus additional 1 hr co-incubation with 10 ng/ml TNFα (or DMEM control) prior to fixation. n = 3 biological replicates. Cell feature measurements were calculated as fold changes to controls (DMSO and BSA/PBS) then Z-scored across TNFα treatments and cytoskeletal drugs by cell line. Nuclear RELA measurements were normalised to the mean of all measurements by cell feature across all TNFα treatments by cell line.

### Quantification and Statistical Analysis

To analyse cell-to-cell differences in the 35 geometric, cytoskeletal and Hoechst features within and between PDAC cell lines, single cell and well (mean) data were collated from all cell lines and treatments (TNFα 0, 0.01, 0.1 and 10 ng/ml) from three biological replicates. Features were normalised to the mean across all treatments and cell lines for each biological replicate. Features were reduced for Bayesian analysis by clustering normalised single cell measurements into ten clusters using the ‘ComplexHeatmap’ package in R (Gu et al., 2016), clustering by the Pearson coefficient with complete linkage, as shown in Figure 2 – Figure supplement 1B. Bayesian network models and arc strengths were generated in R using normalised single cell data for the ten reduced features via the ‘bnlearn’ R package (rsmax2 method) (Scutari, 2010). This algorithm depicts unidirectional arcs, so reverse relationships can exist but are not as statistically likely as the directional relationships indicated. Bayesian modelling with cytoskeletal perturbation used data collated from all small molecules and TNFα doses (0, 0.01, 0.1 and 10 ng/ml) for MIA PaCa2 and PANC1 cells.

Z-scores in Figure 2 – Figure supplement 2 were calculated per technical replicate using the mean and standard deviation for control measurements for each feature across all lines for each biological replicate (N = 3). Mean Z-scores per feature and cell line were calculated by averaging Z-scores for technical replicates across all biological replicates.

Statistical tests were carried out using the ‘Rstatix’ package and visualised using the ‘ggpubr’ package in R. Principal Component Analysis (PCA) was carried out in R using the inbuilt ‘prcomp’ function using Z-score data. Graphs were generated in R using the ‘ggplot’ package (Wickham, 2016).

## Data Availability

All data generated for this study have been included as source data files.

---

**Figure 1 - figure supplement 1:**
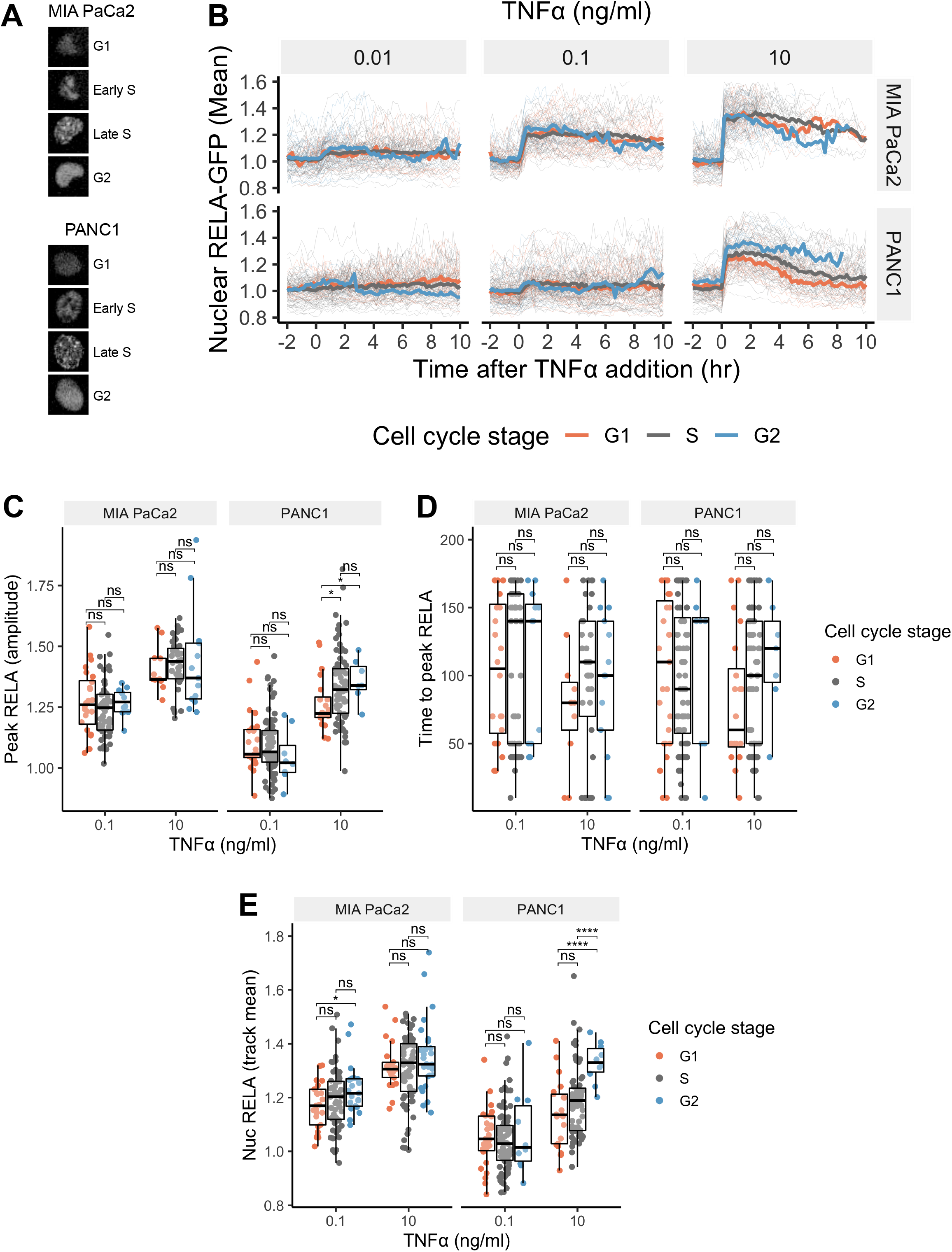
Cell cycle independence of TNFα-mediated RELA dynamics in PDAC cells. (A) Confocal microscopy images of PCNA-Scarlet in representative MIA PaCa2 and PANC1 nuclei in G1, early S, late S or G2 cell cycle stage. (B) Single cell nuclear RELA-GFP tracks for MIA PaCa2 and PANC1 cells categorised by cell cycle stage at the time of TNFα addition (0.01,0.1 or 10 ng/ml), based on the intensity and appearance of PCNA-Scarlet. (C) Single cell measurements of nuclear RELA intensity at track peak and (D) time to track peak by cell cycle stage at TNFα addition. (E) Mean nuclear RELA intensity per single cell track, by cell cycle stage at TNFα addition.

**Figure 2 - figure supplement 1:**
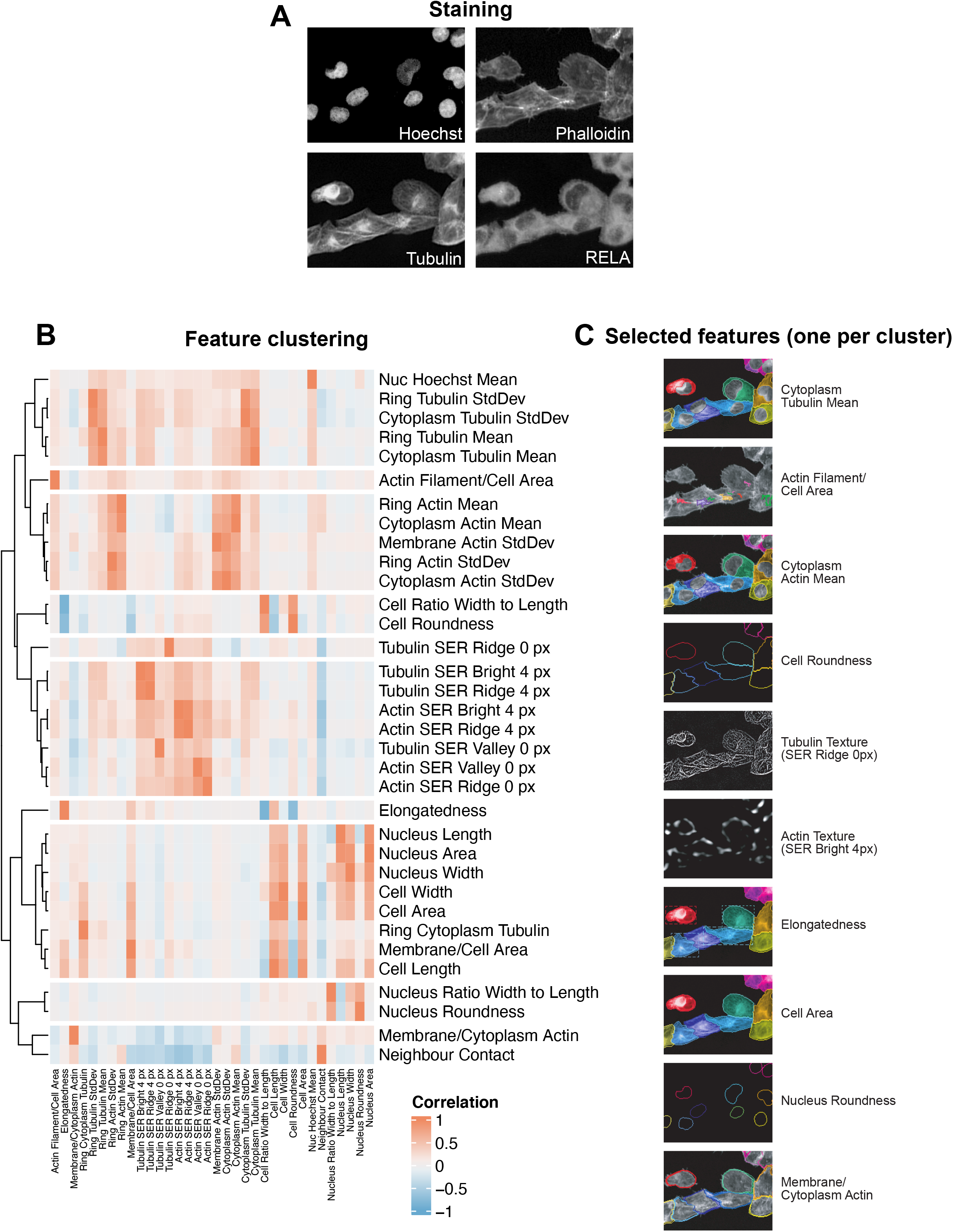
Automated image analysis of RELA localisation, cell shape, and the cytoskeleton in human cells. (A) Staining by immunofluorescence (α-tubulin and RELA) and dyes marking DNA (Hoechst) and F-actin (phalloidin). Cell regions were segmented using Hoechst and α-Tubulin stains and features were measured in these regions using the four stains. (B) Hierarchical clustering of 35 normalised cell features (excluding RELA measurements) measured in five PDAC cell lines. (C) Selected features (one per cluster) are highlighted in the dendrogram and displayed in the images beside.

**Figure 2 - figure supplement 2:**
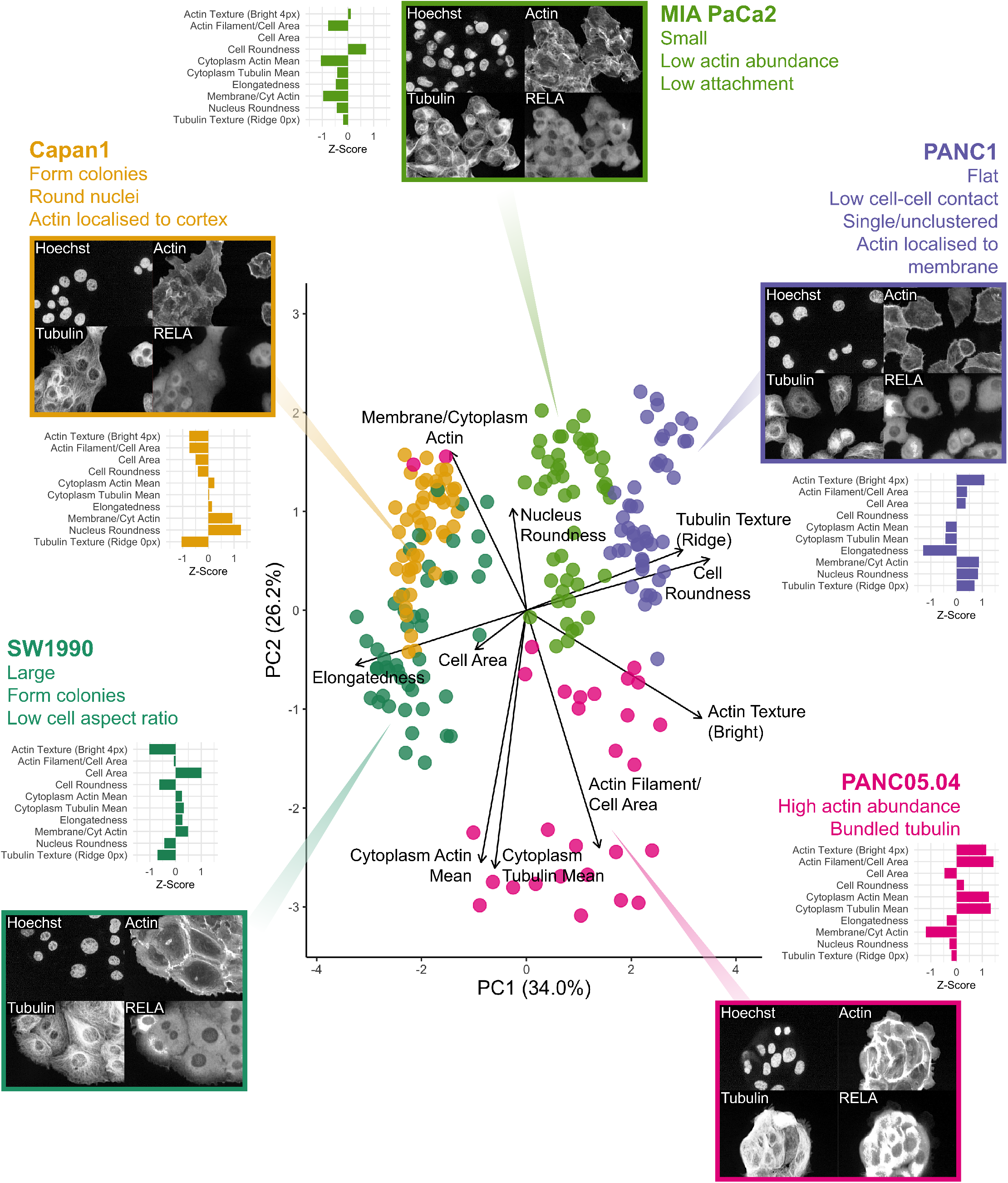
Principal components analysis of five human PDAC cell lines with a reduced set of cytoskeletal and cell shape features. Principal Component Analysis (PCA) of the ten reduced features used in Bayesian analysis for five PDAC cell lines using two principal components (PC1 and PC2). Overlayed is a biplot (arrows) as an indication of the contribution of each cell feature to the two PCs. Circles represent technical replicates. n = four biological repeats. Horizontal bar graphs for individual cell lines show mean z-scores for each feature, calculated across the six cell lines to compare differences in each feature between the cell lines. Z-scores were calculated using the mean and standard deviation for control measurements across all lines for each biological repeat. Displayed is the mean Z score across all biological repeats (n = 4). Images for each cell line show Hoechst, phalloidin marking F-actin (Actin), α-tubulin (Tubulin) and RELA staining by immunofluorescence.

**Figure 2 - figure supplement 3:**
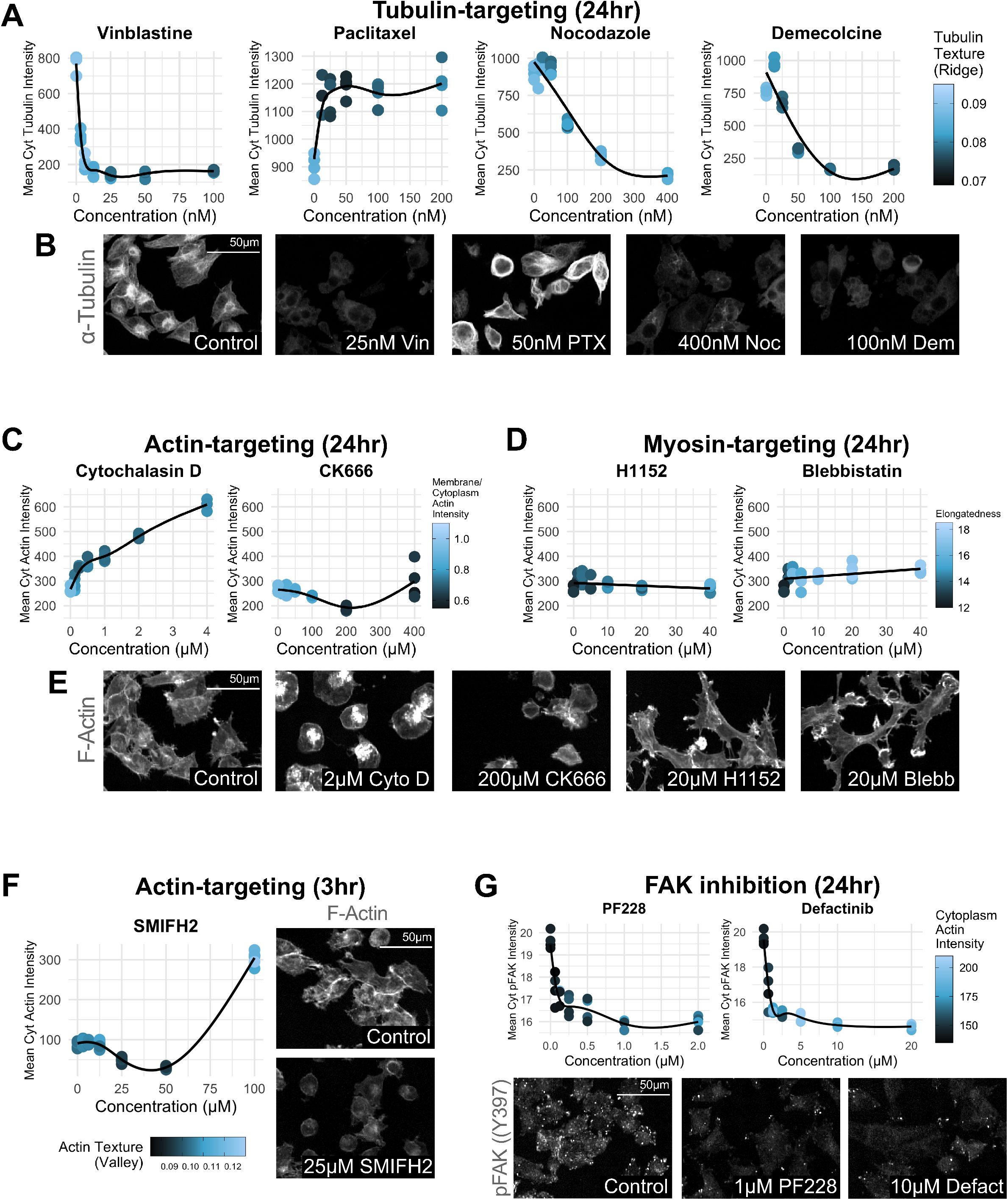
Dose responses for cytoskeleton-targeting drugs. Dose response curves and representative images of MIA PaCa2 cells treated with small molecules in the absence of TNFα. n = four technical replicates per drug dose. (A & B) 24 hr treatment with tubulin-targeting: vinblastine (0-100 nM), paclitaxel (0-200 nM), nocodazole (0-400 nM) and demelcoline (0-200 nM). (C & E) 24 hr treatment with actin-targeting drugs: cytochalasin D (0-4 μM) and CK666 (0-400 μM). (D & E) 24 hr treatment with myosin-targeting drugs: H1152 (0-40 μM) and blebbistatin (0-40 μM). (F) 3 hr treatment with the actin-targeting drug SMIFH2 (0-100 μM). (G) 24 hr treatment with the FAK inhibitors PF-573228 (PF228; 0-2 μM) and defactinib (0-20 μM).

**Figure 3 - figure supplement 1:**
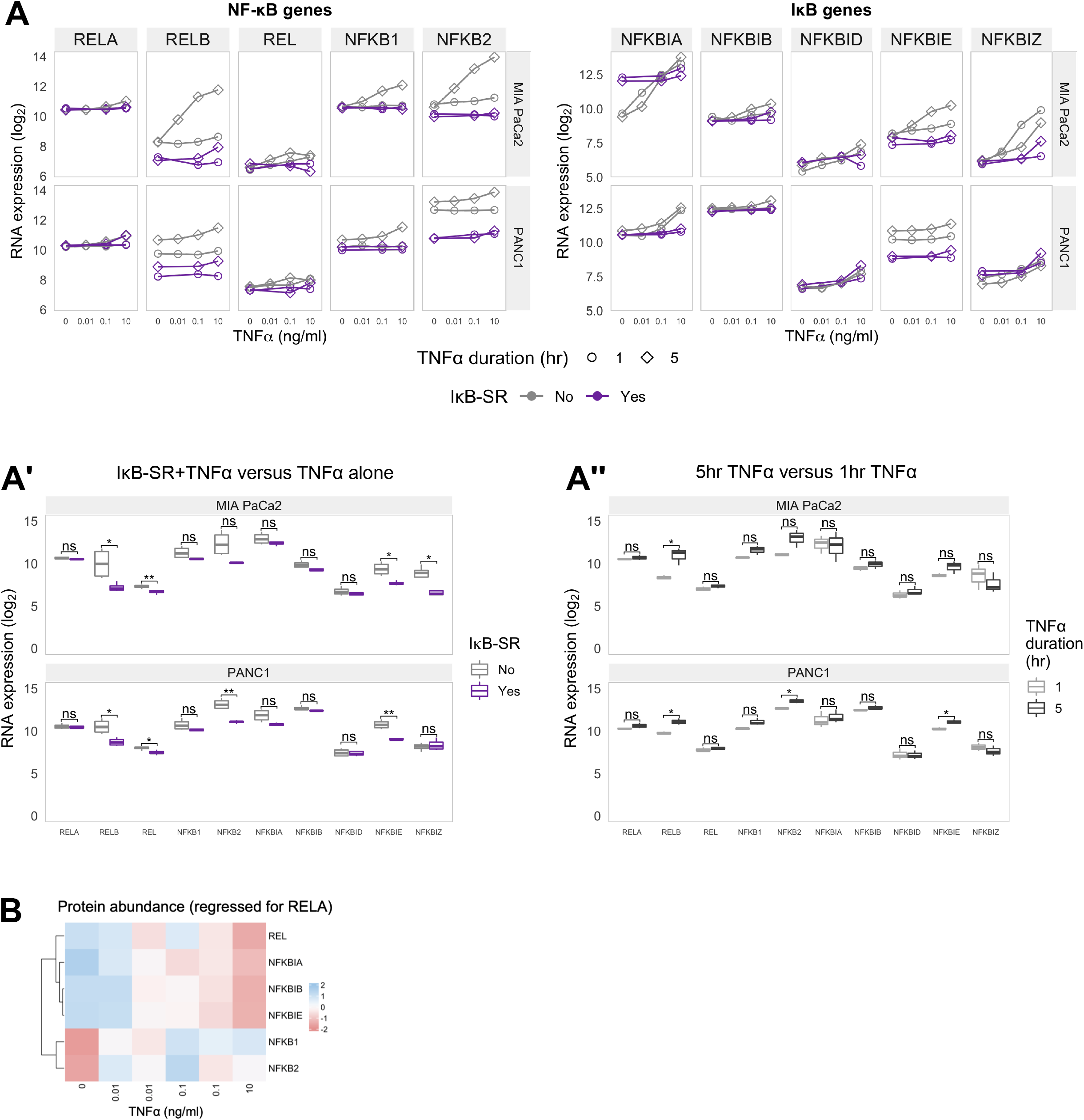
Hierarchical clustering of TNFα/RELA-regulated gene expression in PDAC cells. (A) RNA-seq expression of genes encoding NF-κB (left) and IκB (proteins) in MIA PaCa2 and PANC1 cells treated with TNFα for 1 or 5 hr, in the presence of IκB-SR (constitutively active IκB) or a DMSO control. Normalised counts were generated by DESeq2 and log_2_ transformed. (A” and A”’) Statistical analysis (t-tests with Benjamini-Hochberg correction) of data in (A) comparing RNA expression of each gene (A’’) between cells treated with TNFα (data from 0.1 and 10 ng/ml, and 1 hr and 5 hr treatment collated) with the presence or absence of IkB-SR; or (A’’’) between cells with 1 hr or 5 hr TNFα treatment (0.1 and 10 ng/ml collated) in the absence of IkB-SR. ns (non-significant) = p > 0.05, * = 0 < 0.05, ** = p < 0.01. (B) Summary of proteins detected by Co-IP and mass spectrometry using GFP-Trap in MIA PaCa2 RELA-GFP cells. Cells were treated with TNFα (0, 0.01, 0.1 or 10 ng/ml) for 1 hr. Pull-downs were analysed using mass spectrometry. n = 1 experimental repeats for 0 and 10 ng/ml TNFα and n = 2 experiments repeats for 0.01 and 0.1 ng/ml TNFα. Protein abundances at each TNFα dose adjusted to RELA abundance.

**Figure 4 - figure supplement 1:**
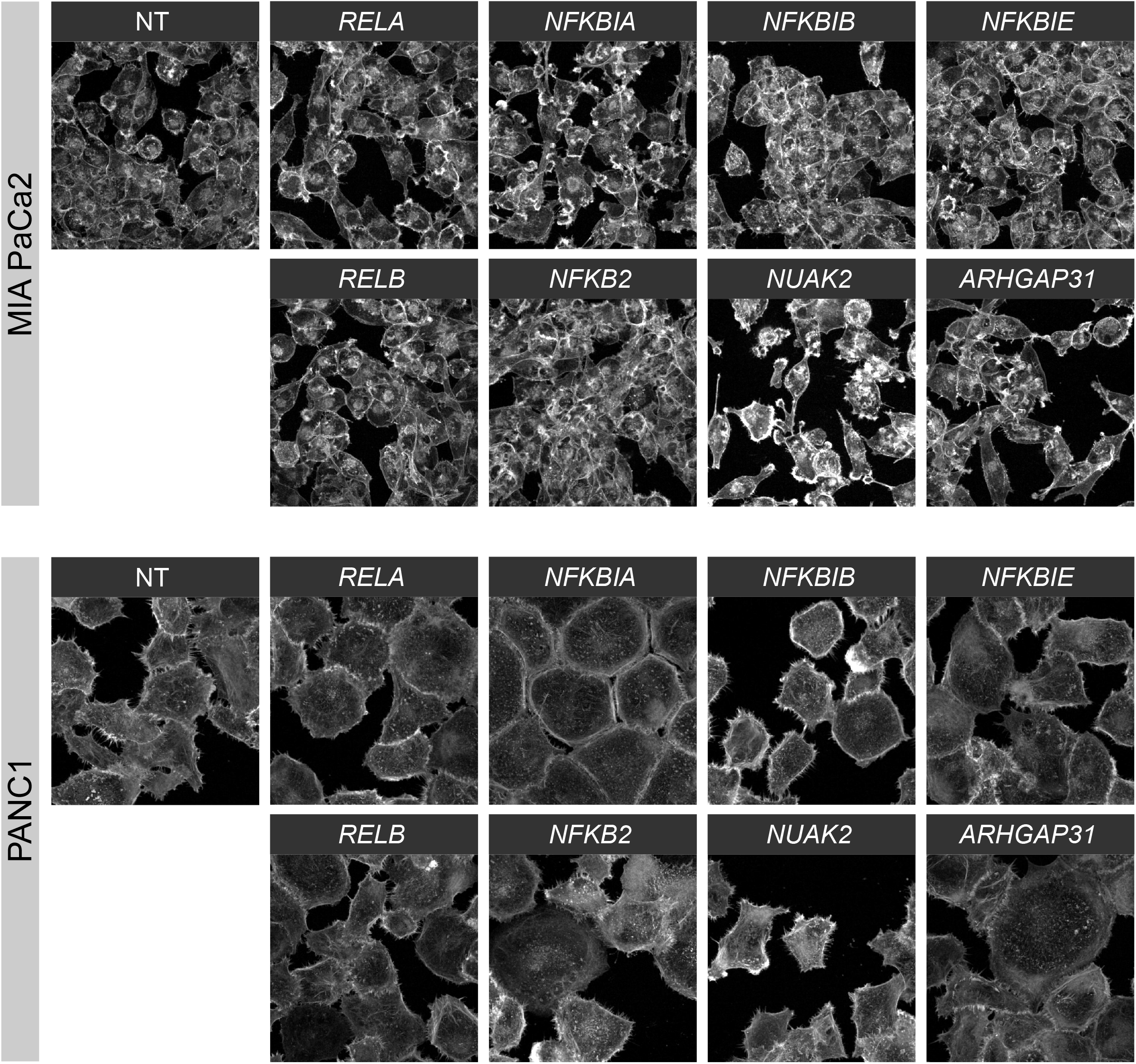
siRNAs targeting the NF-κB pathway and the NF-κB regulated genes *NUAK2* and *ARHGAP31* affect cell morphology and actin organisation. Confocal images of mean actin intensity in the cytoplasm of MIA PaCa2 and PANC1 cells transfected with siRNAs for 48 hr: non-targeting (NT); the NF-κB proteins RELA, RELB, and NFKB2; or the NF-κB inhibitors NFKBIA, NFKBIB, and NFKBIE. Cells were stained with phalloidin for F-actin visualisation.

## References

Ashall, L., Horton, C.A., Nelson, D.E., Paszek, P., Harper, C.V., Sillitoe, K., Ryan, S., Spiller, D.G., Unitt, J.F., Broomhead, D.S., Kell, D.B., Rand, D.A., Sée, V., White, M.R.H., 2009. Pulsatile stimulation determines timing and specificity of NF-κB-dependent transcription. Science 324, 242–246. https://doi.org/10.1126/science.1164860

Baeuerle, P.A., Baltimore, D., 1988. Activation of DNA-binding activity in an apparently cytoplasmic precursor of the NF-kappa B transcription factor. Cell 53, 211–217. https://doi.org/10.1016/0092-8674(88)90382-0

Barr, A.R., Cooper, S., Heldt, F.S., Butera, F., Stoy, H., Mansfeld, J., Novák, B., Bakal, C., 2017. DNA damage during S-phase mediates the proliferation-quiescence decision in the subsequent G1 via p21 expression. Nat. Commun. 8, 1–17. https://doi.org/10.1038/ncomms14728

Bengtsson, A., Andersson, R., Ansari, D., 2020. The actual 5-year survivors of pancreatic ductal adenocarcinoma based on real-world data. Sci. Rep. 10, 16425. https://doi.org/10.1038/s41598-020-73525-y

Bold, R.J., Virudachalam, S., McConkey, D.J., 2001. Chemosensitization of Pancreatic Cancer by Inhibition of the 26S Proteasome. J. Surg. Res. 100, 11–17. https://doi.org/10.1006/jsre.2001.6194

Bourgarel-Rey, V., Vallee, S., Rimet, O., Champion, S., Braguer, D., Desobry, A., Briand, C., Barra, Y., 2001. Involvement of nuclear factor kappaB in c-Myc induction by tubulin polymerization inhibitors. Mol. Pharmacol. 59, 1165–1170. https://doi.org/10.1124/mol.59.5.1165

Bren, G.D., Solan, N.J., Miyoshi, H., Pennington, K.N., Pobst, L.J., Paya, C.V., 2001. Transcription of the RelB gene is regulated by NF-kappaB. Oncogene 20, 7722–7733. https://doi.org/10.1038/sj.onc.1204868

Brown, K., Park, S., Kanno, T., Franzoso, G., Siebenlist, U., 1993. Mutual regulation of the transcriptional activator NF-kappa B and its inhibitor, I kappa B-alpha. Proc. Natl. Acad. Sci. U. S. A. 90, 2532–2536. https://doi.org/10.1073/pnas.90.6.2532

Chen, Z., Hagler, J., Palombella, V.J., Melandri, F., Scherer, D., Ballard, D., Maniatis, T., 1995. Signal-induced site-specific phosphorylation targets I kappa B alpha to the ubiquitin-proteasome pathway. Genes Dev. 9, 1586–1597. https://doi.org/10.1101/gad.9.13.1586

Collart, M.A., Baeuerle, P., Vassalli, P., 1990. Regulation of tumor necrosis factor alpha transcription in macrophages: involvement of four kappa B-like motifs and of constitutive and inducible forms of NF-kappa B. Mol. Cell. Biol. 10, 1498–1506. https://doi.org/10.1128/mcb.10.4.1498-1506.1990

Deer, E.L., Gonzalez-Hernandez, J., Coursen, J.D., Shea, J.E., Ngatia, J., Scaife, C.L., Firpo, M.A., Mulvihill, S.J., 2010. Phenotype and Genotype of Pancreatic Cancer Cell Lines. Pancreas 39, 425–435. https://doi.org/10.1097/MPA.0b013e3181c15963

DiDonato, J.A., Hayakawa, M., Rothwarf, D.M., Zandi, E., Karin, M., 1997. A cytokine-responsive IκB kinase that activates the transcription factor NF-κB. Nature 388, 548–554. https://doi.org/10.1038/41493

Frasor, J., Weaver, A., Pradhan, M., Dai, Y., Miller, L.D., Lin, C.-Y., Stanculescu, A., 2009. Positive cross-talk between estrogen receptor and NF-kappaB in breast cancer. Cancer Res. 69, 8918–8925. https://doi.org/10.1158/0008-5472.CAN-09-2608

Fujioka, S., Sclabas, G.M., Schmidt, C., Frederick, W.A., Dong, Q.G., Abbruzzese, J.L., Evans, D.B., Baker, C., Chiao, P.J., 2003. Function of Nuclear Factor κB in Pancreatic Cancer Metastasis. Clin. Cancer Res. 9, 346–354.

Ghosh, S., May, M.J., Kopp, E.B., 1998. NF-κB and Rel Proteins: Evolutionarily Conserved Mediators of Immune Responses. Annu. Rev. Immunol. 16, 225–260. https://doi.org/10.1146/annurev.immunol.16.1.225

Gigant, B., Wang, C., Ravelli, R.B.G., Roussi, F., Steinmetz, M.O., Curmi, P.A., Sobel, A., Knossow, M., 2005. Structural basis for the regulation of tubulin by vinblastine. Nature 435, 519–522. https://doi.org/10.1038/nature03566

Hayden, M.S., West, A.P., Ghosh, S., 2006. NF-κB and the immune response. Oncogene 25, 6758–6780. https://doi.org/10.1038/sj.onc.1209943

Heid, I., Lubeseder-Martellato, C., Sipos, B., Mazur, P.K., Lesina, M., Schmid, R.M., Siveke, J.T., 2011. Early requirement of Rac1 in a mouse model of pancreatic cancer. Gastroenterology 141, 719–730, 730.e1-7. https://doi.org/10.1053/j.gastro.2011.04.043

Hoffmann, A., Levchenko, A., Scott, M.L., Baltimore, D., 2002. The IκB-NF-κB Signaling Module: Temporal Control and Selective Gene Activation. Science 298, 1241–1245. https://doi.org/10.1126/science.1071914

Isogai, T., van der Kammen, R., Innocenti, M., 2015. SMIFH2 has effects on Formins and p53 that perturb the cell cytoskeleton. Sci. Rep. 5, 9802. https://doi.org/10.1038/srep09802

Kieran, M., Blank, V., Logeat, F., Vandekerckhove, J., Lottspeich, F., Le Bail, O., Urban, M.B., Kourilsky, P., Baeuerle, P.A., Israël, A., 1990. The DNA binding subunit of NF-kappa B is identical to factor KBF1 and homologous to the rel oncogene product. Cell 62, 1007–1018. https://doi.org/10.1016/0092-8674(90)90275-j

Kovács, M., Tóth, J., Hetényi, C., Málnási-Csizmadia, A., Sellers, J.R., 2004. Mechanism of Blebbistatin Inhibition of Myosin II. J. Biol. Chem. 279, 35557–35563. https://doi.org/10.1074/jbc.M405319200

Kunnumakkara, A.B., Guha, S., Krishnan, S., Diagaradjane, P., Gelovani, J., Aggarwal, B.B., 2007. Curcumin Potentiates Antitumor Activity of Gemcitabine in an Orthotopic Model of Pancreatic Cancer through Suppression of Proliferation, Angiogenesis, and Inhibition of Nuclear Factor-κB–Regulated Gene Products. Cancer Res. 67, 3853–3861. https://doi.org/10.1158/0008-5472.CAN-06-4257

Kurki, P., Vanderlaan, M., Dolbeare, F., Gray, J., Tan, E.M., 1986. Expression of proliferating cell nuclear antigen (PCNA)/cyclin during the cell cycle. Exp. Cell Res. 166, 209–219. https://doi.org/10.1016/0014-4827(86)90520-3

Lamarche-Vane, N., Hall, A., 1998. CdGAP, a novel proline-rich GTPase-activating protein for Cdc42 and Rac. J. Biol. Chem. 273, 29172–29177. https://doi.org/10.1074/jbc.273.44.29172

Lamba, V., Ghosh, I., 2012. New directions in targeting protein kinases: focusing upon true allosteric and bivalent inhibitors. Curr. Pharm. Des. 18, 2936–2945. https://doi.org/10.2174/138161212800672813

Lane, K., Valen, D.V., DeFelice, M.M., Macklin, D.N., Kudo, T., Jaimovich, A., Carr, A., Meyer, T., Pe’er, D., Boutet, S.C., Covert, M.W., 2017. Measuring Signaling and RNA-Seq in the Same Cell Links Gene Expression to Dynamic Patterns of NF-κB Activation. Cell Syst. 4, 458–469.e5. https://doi.org/10.1016/j.cels.2017.03.010

Libermann, T.A., Baltimore, D., 1990. Activation of interleukin-6 gene expression through the NF-kappa B transcription factor. Mol. Cell. Biol. 10, 2327–2334. https://doi.org/10.1128/mcb.10.5.2327-2334.1990

Lombardi, L., Ciana, P., Cappellini, C., Trecca, D., Guerrini, L., Migliazza, A., Maiolo, A.T., Neri, A., 1995. Structural and functional characterization of the promoter regions of the NFKB2 gene. Nucleic Acids Res. 23, 2328–2336. https://doi.org/10.1093/nar/23.12.2328

Melisi, D., Niu, J., Chang, Z., Xia, Q., Peng, B., Ishiyama, S., Evans, D.B., Chiao, P.J., 2009. Secreted interleukin-1alpha induces a metastatic phenotype in pancreatic cancer by sustaining a constitutive activation of nuclear factor-kappaB. Mol. Cancer Res. MCR 7, 624–633. https://doi.org/10.1158/1541-7786.MCR-08-0201

Mullins, R.D., Heuser, J.A., Pollard, T.D., 1998. The interaction of Arp2/3 complex with actin: Nucleation, high affinity pointed end capping, and formation of branching networks of filaments. Proc. Natl. Acad. Sci. 95, 6181–6186. https://doi.org/10.1073/pnas.95.11.6181

Nelson, D.E., Ihekwaba, A.E.C., Elliott, M., Johnson, J.R., Gibney, C.A., Foreman, B.E., Nelson, G., See, V., Horton, C.A., Spiller, D.G., Edwards, S.W., McDowell, H.P., Unitt, J.F., Sullivan, E., Grimley, R., Benson, N., Broomhead, D., Kell, D.B., White, M.R.H., 2004. Oscillations in NF-κB Signaling Control the Dynamics of Gene Expression. Science 306, 704–708. https://doi.org/10.1126/science.1099962

Németh, Z.H., Deitch, E.A., Davidson, M.T., Szabó, C., Vizi, E.S., Haskó, G., 2004. Disruption of the actin cytoskeleton results in nuclear factor-kappaB activation and inflammatory mediator production in cultured human intestinal epithelial cells. J. Cell. Physiol. 200, 71–81. https://doi.org/10.1002/jcp.10477

Nolan, G.P., Ghosh, S., Liou, H.C., Tempst, P., Baltimore, D., 1991. DNA binding and I kappa B inhibition of the cloned p65 subunit of NF-kappa B, a rel-related polypeptide. Cell 64, 961–969. https://doi.org/10.1016/0092-8674(91)90320-x

Pe’er, D., Regev, A., Elidan, G., Friedman, N., 2001. Inferring subnetworks from perturbed expression profiles. Bioinforma. Oxf. Engl. 17 Suppl 1, S215–224. https://doi.org/10.1093/bioinformatics/17.suppl_1.s215

Pruyne, D., Evangelista, M., Yang, C., Bi, E., Zigmond, S., Bretscher, A., Boone, C., 2002. Role of formins in actin assembly: nucleation and barbed-end association. Science 297, 612–615. https://doi.org/10.1126/science.1072309

Razidlo, G.L., Burton, K.M., McNiven, M.A., 2018. Interleukin-6 promotes pancreatic cancer cell migration by rapidly activating the small GTPase CDC42. J. Biol. Chem. 293, 11143–11153. https://doi.org/10.1074/jbc.RA118.003276

Rizvi, S.A., Neidt, E.M., Cui, J., Feiger, Z., Skau, C.T., Gardel, M.L., Kozmin, S.A., Kovar, D.R., 2009. Identification and Characterization of a Small Molecule Inhibitor of Formin-Mediated Actin Assembly. Chem. Biol. 16, 1158–1168. https://doi.org/10.1016/j.chembiol.2009.10.006

Rosette, C., Karin, M., 1995. Cytoskeletal control of gene expression: depolymerization of microtubules activates NF-kappa B. J. Cell Biol. 128, 1111–1119.

Sachs, K., Perez, O., Pe’er, D., Lauffenburger, D.A., Nolan, G.P., 2005. Causal Protein-Signaling Networks Derived from Multiparameter Single-Cell Data. Science 308, 523–529. https://doi.org/10.1126/science.1105809

Sailem, H.Z., Bakal, C., 2017. Identification of clinically predictive metagenes that encode components of a network coupling cell shape to transcription by image-omics. Genome Res. 27, 196–207. https://doi.org/10.1101/gr.202028.115

Sasaki, Y., Suzuki, M., Hidaka, H., 2002. The novel and specific Rho-kinase inhibitor (S)-(+)-2-methyl-1-[(4-methyl-5-isoquinoline)sulfonyl]-homopiperazine as a probing molecule for Rho-kinase-involved pathway. Pharmacol. Ther. 93, 225–232.

Scott, M.L., Fujita, T., Liou, H.C., Nolan, G.P., Baltimore, D., 1993. The p65 subunit of NF-kappa B regulates I kappa B by two distinct mechanisms. Genes Dev. 7, 1266–1276. https://doi.org/10.1101/gad.7.7a.1266

Scutari, M., Graafland, C.E., Gutierrez, J.M., 2018. Who Learns Better Bayesian Network Structures: Constraint-Based, Score-based or Hybrid Algorithms? Proc. Mach. Learn. Res. 72, 416–427.

Sero, J.E., Sailem, H.Z., Ardy, R.C., Almuttaqi, H., Zhang, T., Bakal, C., 2015. Cell shape and the microenvironment regulate nuclear translocation of NF-κB in breast epithelial and tumor cells. Mol. Syst. Biol. 11. https://doi.org/10.15252/msb.20145644

Spencer, C.M., Faulds, D., 1994. Paclitaxel. Drugs 48, 794–847. https://doi.org/10.2165/00003495-199448050-00009

Stewart-Ornstein, J., Lahav, G., 2016. Dynamics of CDKN1A in Single Cells Defined by an Endogenous Fluorescent Tagging Toolkit. Cell Rep. 14, 1800–1811. https://doi.org/10.1016/j.celrep.2016.01.045

Sun, S.C., Ganchi, P.A., Ballard, D.W., Greene, W.C., 1993. NF-kappa B controls expression of inhibitor I kappa B alpha: evidence for an inducible autoregulatory pathway. Science 259, 1912–1915. https://doi.org/10.1126/science.8096091

Sung, M.-H., Salvatore, L., Lorenzi, R.D., Indrawan, A., Pasparakis, M., Hager, G.L., Bianchi, M.E., Agresti, A., 2009. Sustained Oscillations of NF-κB Produce Distinct Genome Scanning and Gene Expression Profiles. PLOS ONE 4, e7163. https://doi.org/10.1371/journal.pone.0007163

Tangutur, A.D., Kumar, D., Krishna, K.V., Kantevari, S., 2017. Microtubule Targeting Agents as Cancer Chemotherapeutics: An Overview of Molecular Hybrids as Stabilizing and Destabilizing Agents. Curr. Top. Med. Chem. 17, 2523–2537. https://doi.org/10.2174/1568026617666170104145640

Tay, S., Hughey, J.J., Lee, T.K., Lipniacki, T., Quake, S.R., Covert, M.W., 2010. Single-cell NF-κB dynamics reveal digital activation and analog information processing in cells. Nature 466, 267–271. https://doi.org/10.1038/nature09145

Tcherkezian, J., Triki, I., Stenne, R., Danek, E.I., Lamarche-Vane, N., 2006. The human orthologue of CdGAP is a phosphoprotein and a GTPase-activating protein for Cdc42 and Rac1 but not RhoA. Biol. Cell 98, 445–456. https://doi.org/10.1042/BC20050101

Vallenius, T., Vaahtomeri, K., Kovac, B., Osiceanu, A.-M., Viljanen, M., Mäkelä, T.P., 2011. An association between NUAK2 and MRIP reveals a novel mechanism for regulation of actin stress fibers. J. Cell Sci. 124, 384–393. https://doi.org/10.1242/jcs.072660

Vasquez, R.J., Howell, B., Yvon, A.M., Wadsworth, P., Cassimeris, L., 1997. Nanomolar concentrations of nocodazole alter microtubule dynamic instability in vivo and in vitro. Mol. Biol. Cell 8, 973–985. https://doi.org/10.1091/mbc.8.6.973

Wang, W., Abbruzzese, J.L., Evans, D.B., Larry, L., Cleary, K.R., Chiao, P.J., 1999. The Nuclear Factor-κB RelA Transcription Factor Is Constitutively Activated in Human Pancreatic Adenocarcinoma Cells. Clin. Cancer Res. 5, 119–127.

Weichert, W., Boehm, M., Gekeler, V., Bahra, M., Langrehr, J., Neuhaus, P., Denkert, C., Imre, G., Weller, C., Hofmann, H.-P., Niesporek, S., Jacob, J., Dietel, M., Scheidereit, C., Kristiansen, G., 2007. High expression of RelA/p65 is associated with activation of nuclear factor-*κ*B-dependent signaling in pancreatic cancer and marks a patient population with poor prognosis. Br. J. Cancer 97, 523–530. https://doi.org/10.1038/sj.bjc.6603878

Yamamoto, H., Takashima, S., Shintani, Y., Yamazaki, S., Seguchi, O., Nakano, A., Higo, S., Kato, H., Liao, Y., Asano, Y., Minamino, T., Matsumura, Y., Takeda, H., Kitakaze, M., 2008. Identification of a novel substrate for TNFalpha-induced kinase NUAK2. Biochem. Biophys. Res. Commun. 365, 541–547. https://doi.org/10.1016/j.bbrc.2007.11.013

Yuan, W.-C., Pepe-Mooney, B., Galli, G.G., Dill, M.T., Huang, H.-T., Hao, M., Wang, Y., Liang, H., Calogero, R.A., Camargo, F.D., 2018. NUAK2 is a critical YAP target in liver cancer. Nat. Commun. 9, 4834. https://doi.org/10.1038/s41467-018-07394-5

Zambrano, S., De Toma, I., Piffer, A., Bianchi, M.E., Agresti, A., 2016. NF-κB oscillations translate into functionally related patterns of gene expression. eLife 5. https://doi.org/10.7554/eLife.09100

Zandi, E., Rothwarf, D.M., Delhase, M., Hayakawa, M., Karin, M., 1997. The IkappaB kinase complex (IKK) contains two kinase subunits, IKKalpha and IKKbeta, necessary for IkappaB phosphorylation and NF-kappaB activation. Cell 91, 243–252. https://doi.org/10.1016/s0092-8674(00)80406-7

Zhao, X., Fan, W., Xu, Z., Chen, H., He, Y., Yang, Gui, Yang, Gang, Hu, H., Tang, S., Wang, P., Zhang, Z., Xu, P., Yu, M., 2016. Inhibiting tumor necrosis factor-alpha diminishes desmoplasia and inflammation to overcome chemoresistance in pancreatic ductal adenocarcinoma. Oncotarget 7, 81110–81122. https://doi.org/10.18632/oncotarget.13212

